## Supplementary material for "Multiple lineages of *Streptomyces* produce antimicrobials within passalid beetle galleries across eastern North America": Figure Supplements

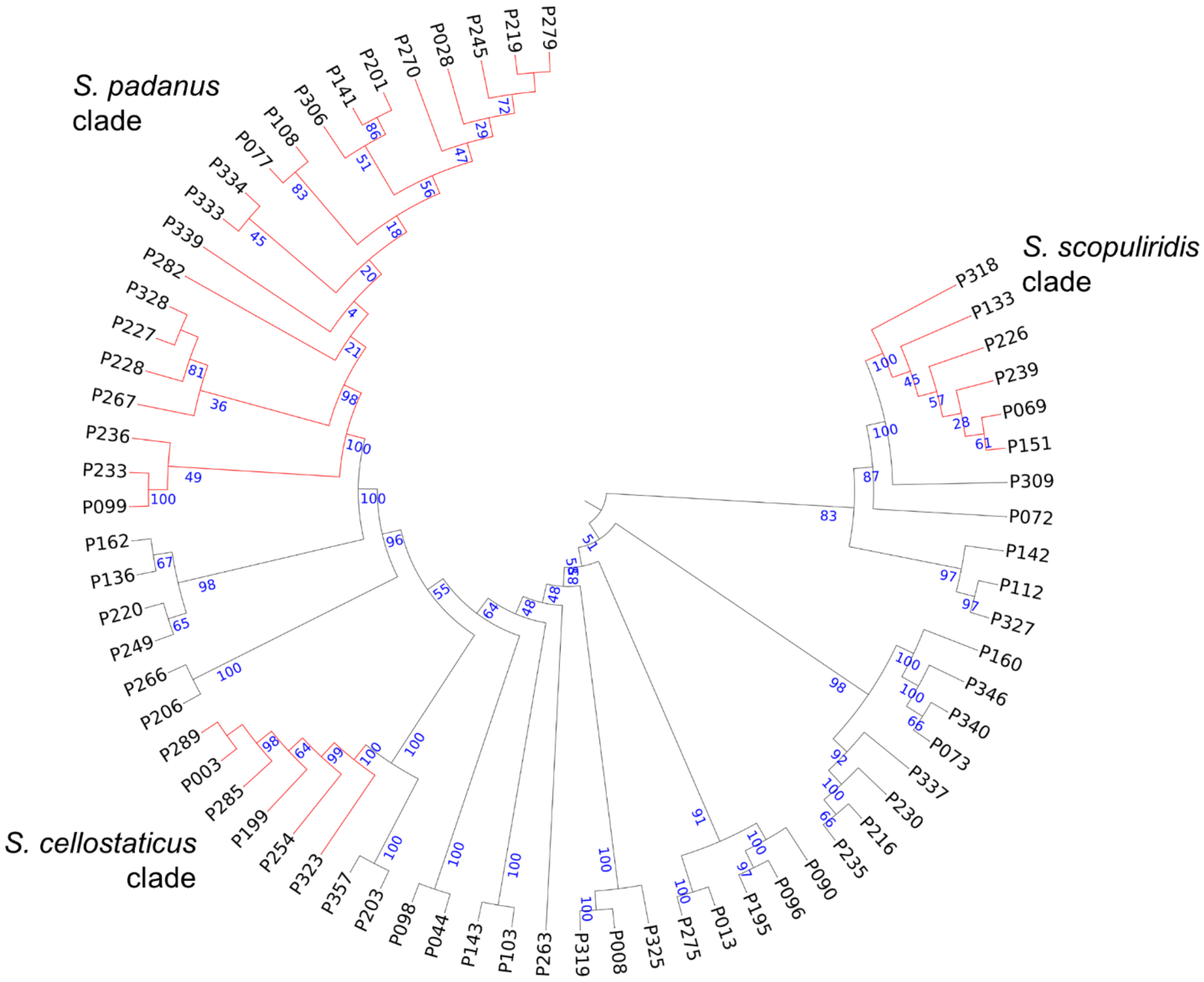


**Figure 4 - figure supplement 1:** Maximum-likelihood phylogenetic tree built using concatenated sequences of four genes (16S rRNA, rpoB, gyrB, atpD). Bootstrap support values (in percentage) are based on 1,000 replicates (numbers in blue). Branches in red highlight the three major clades: *S. padanus*, *S. cellostaticus* and *S. scopuliridis*. Leaf labels represent the strain code. The outgroup (*Mycobacterium tuberculosis* H37RV) was manually removed and the branch length information was not incorporated into the tree to facilitate visualization of the bootstrap values.

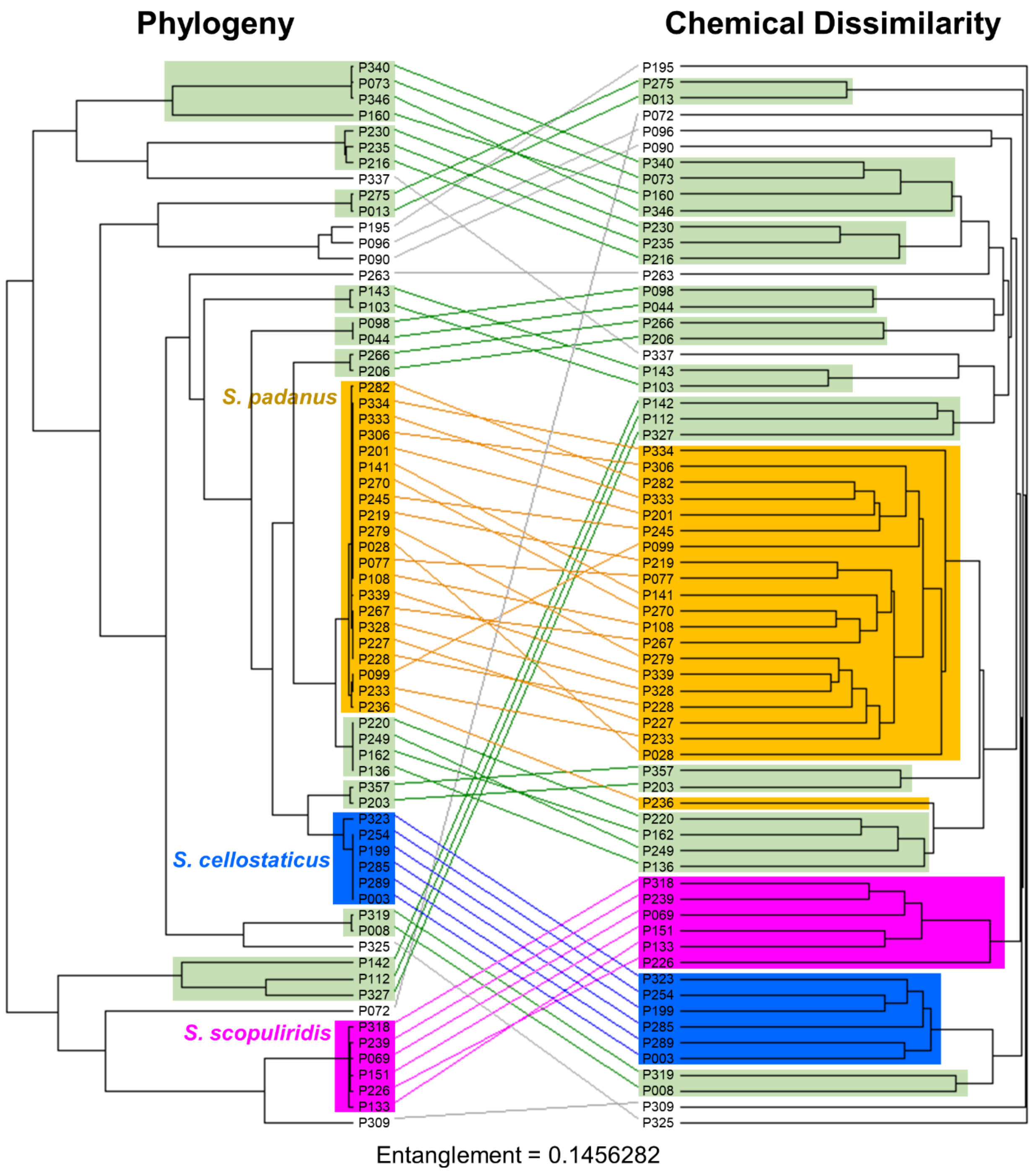


**Figure 4 - figure supplement 2:** Tanglegram analysis comparing phylogenetic (left) and metabolic (right) information of *Streptomyces* strains associated with *O. disjunctus* shows chemo-evolutionary relationships among strains. The maximum-likelihood phylogenetic tree was built using concatenated sequences of four genes (16S rRNA, rpoB, gyrB, atpD). The chemical dissimilarity dendrogram was generated using hierarchical cluster analysis on the presence and absence of ~19,000 chemical features detected in an untargeted metabolomics analysis of culture extracts, using Jaccard distance and UPGMA as the agglomeration method. Lines connects the same strains; orange boxes highlight the *S. padanus* clade; blue boxes highlight the *S. cellostaticus* clade; pink boxes highlight the *S. scopuliridis* clade; green boxes highlight smaller clades that were seen in both sides of the tanglegram.

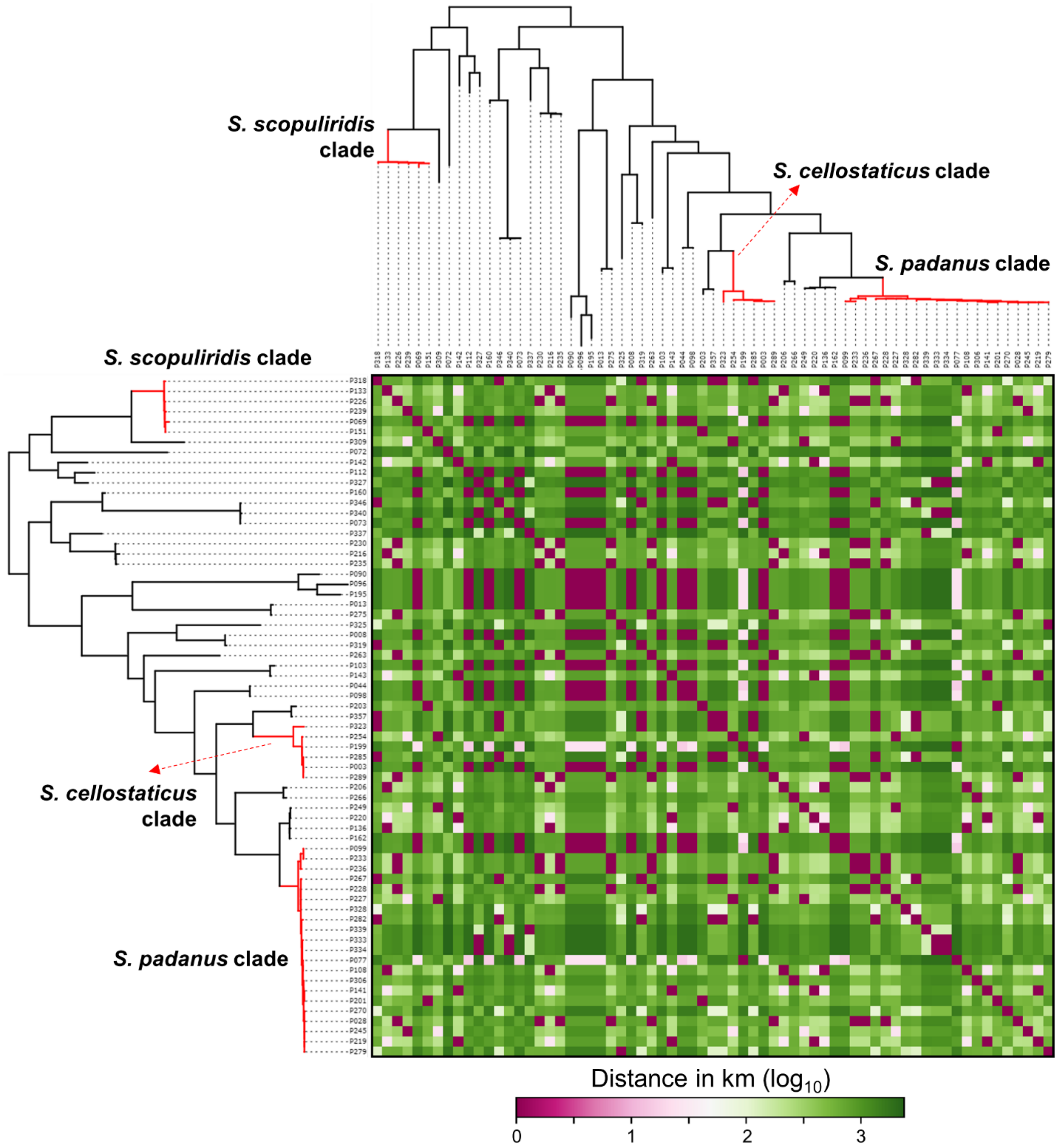


**Figure 4 - figure supplement 3:** Heatmap showing the distance in kilometers (in $log_{10}$ scale) between the geographical origin of *Streptomyces* strains associated with *O. disjunctus* galleries. Branches in red highlight the three major clades: *S. padanus*, *S. cellostaticus* and *S. scopuliridis*. Leaf labels represent the strain code.

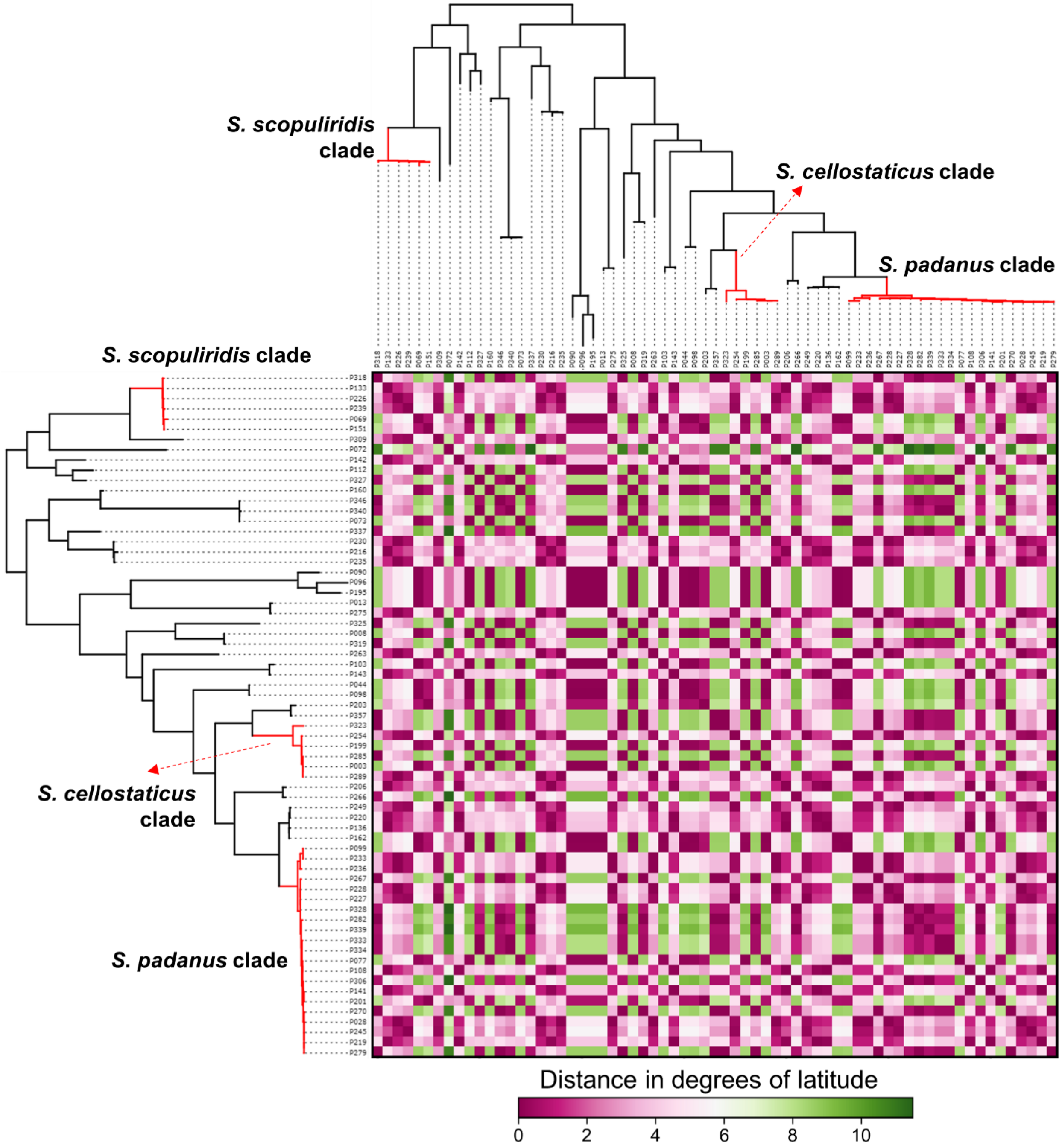


**Figure 4 - figure supplement 4:** Heatmap showing the distance in degrees of latitude between the geographical origin of *Streptomyces* strains associated with *O. disjunctus* galleries. Branches in red highlight the three major clades: *S. padanus*, *S. cellostaticus* and *S. scopuliridis*. Leaf labels represent the strain code.

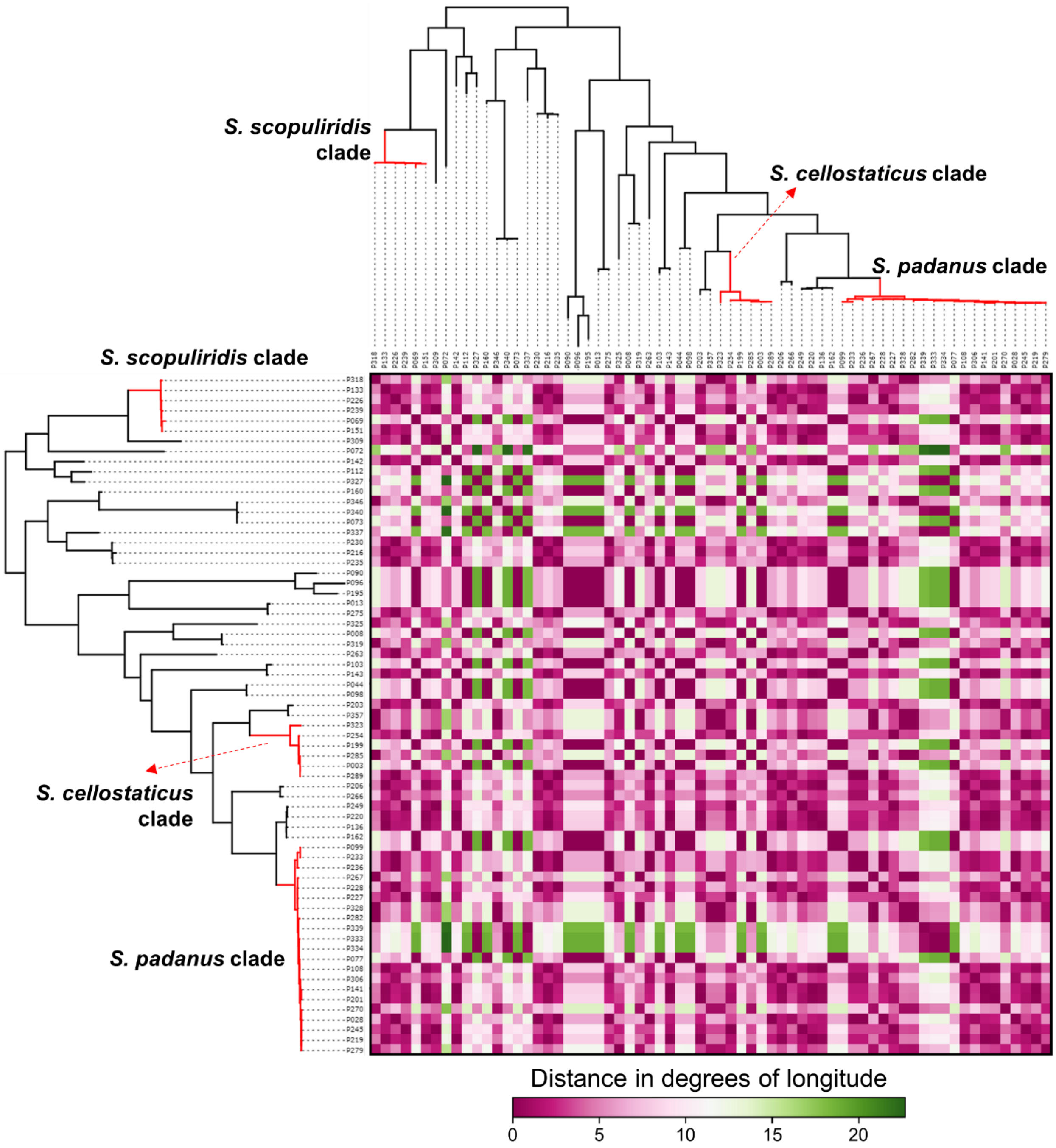


**Figure 4 - figure supplement 5:** Heatmap showing the distance in degrees of longitude between the geographical origin of *Streptomyces* strains associated with *O. disjunctus* galleries. Branches in red highlight the three major clades: *S. padanus*, *S. cellostaticus* and *S. scopuliridis*. Leaf labels represent the strain code.

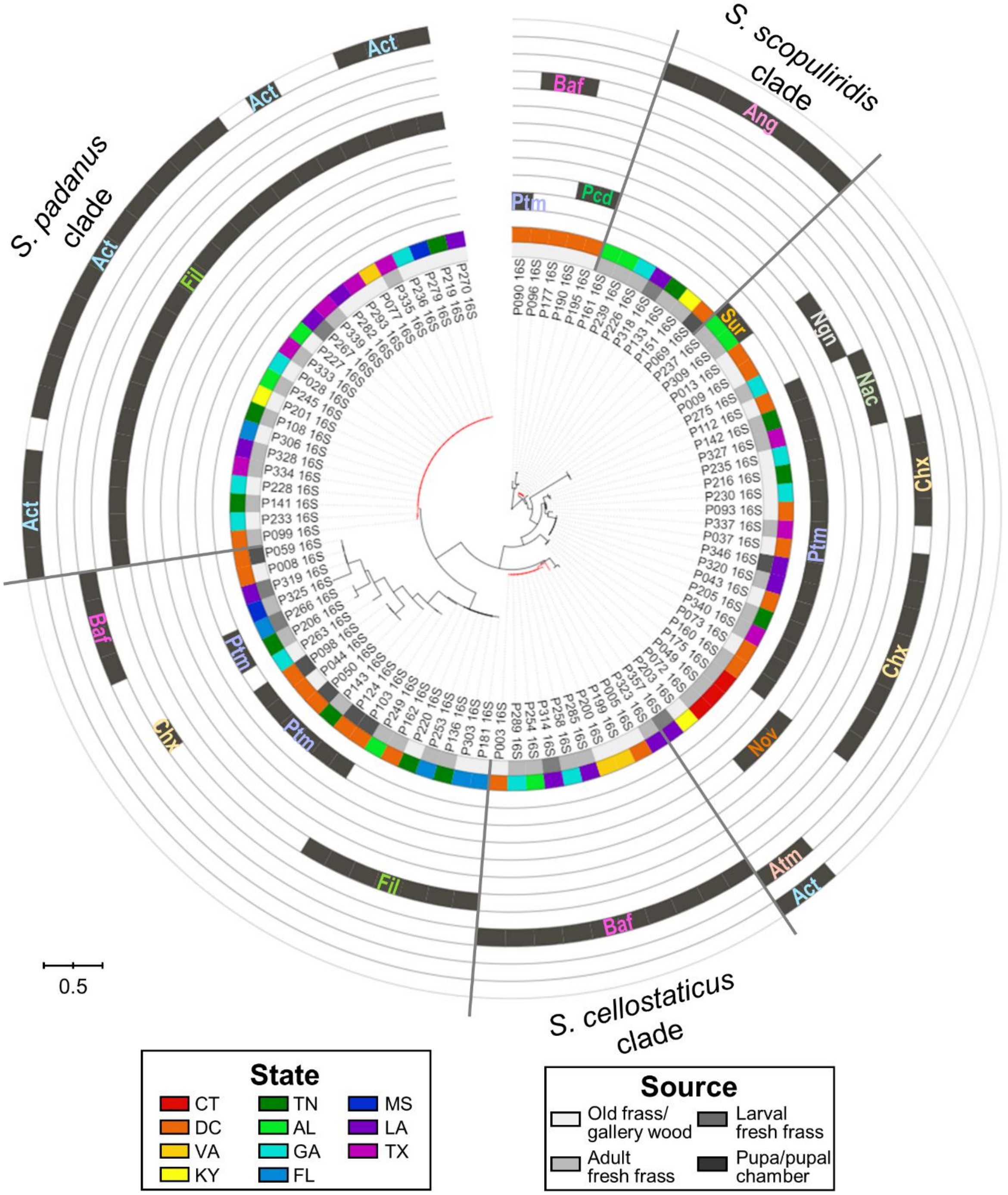


**Figure 4 - figure supplement 6:** Maximum-likelihood phylogenetic tree built using the 16S rRNA gene sequence including duplicated strains, annotated with compounds produced by each microbial strain and their geographic and source origin (both represented by rings around the tree). Scale bar represents branch length in number of substitutions per site. The outgroup (*Mycobacterium tuberculosis* H37RV) was removed manually from the tree to facilitate visualization. Leaf labels represent the strain code. Branches in red highlight the three major clades: *S. padanus*, *S. cellostaticus* and *S. scopuliridis*. Leaf labels represent the strain code. Act: actinomycins. Ang: angucylinones. Atm: antimycins. Baf: bafilomycins. Chx: cycloheximide. Fil: filipins. Nac: nactins. Ngn: nigericin. Nov: novobiocin. Pcd: Piericidin. Ptm: polycyclic tetramate macrolactams. Sur: Surugamides.

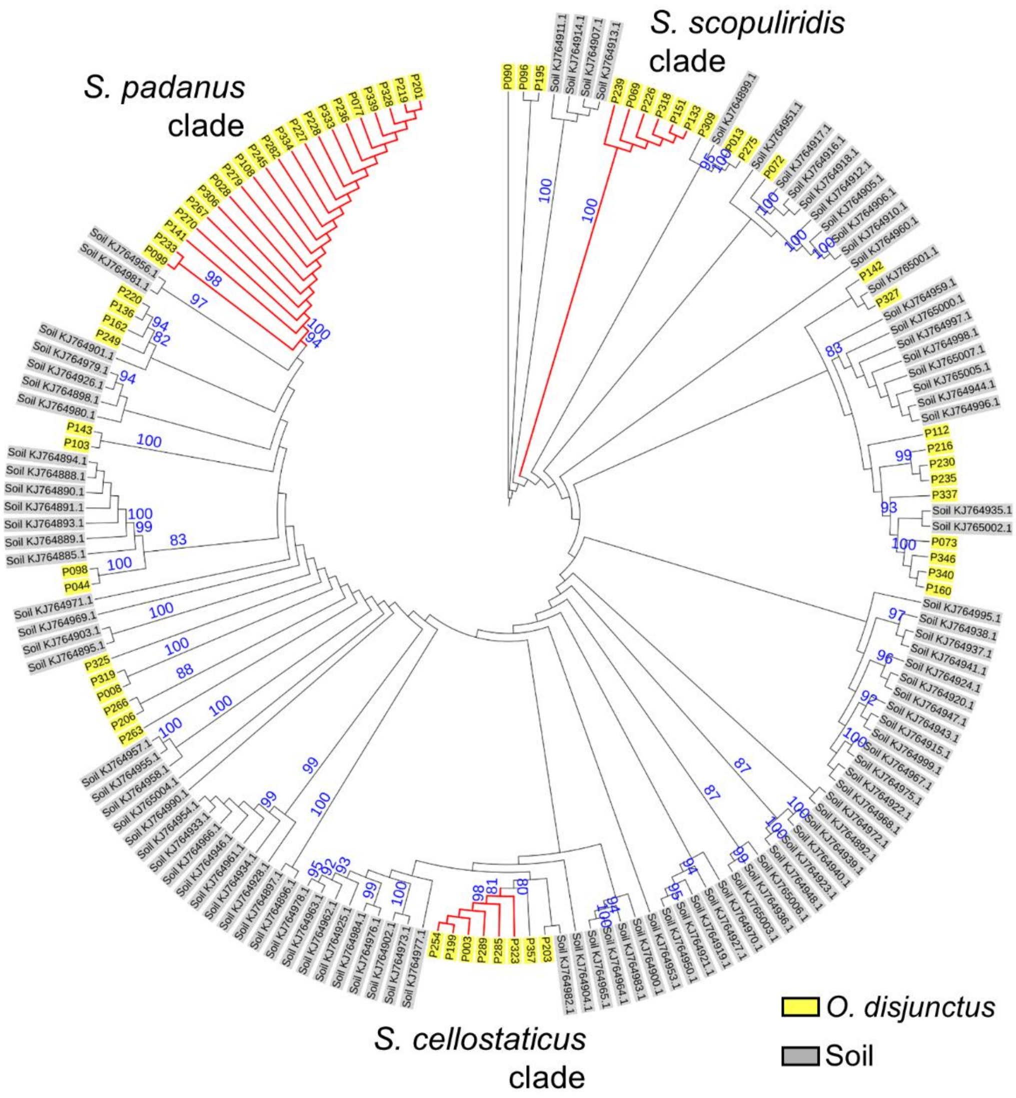


**Figure 4 - figure supplement 7:** Maximum-likelihood phylogenetic tree built using the 16S rRNA gene sequence of *Streptomyces* strains isolated from *O. disjunctus* and soil. Bootstrap support values (in percentage) are based on 1,000 replicates (numbers in blue, only values above 80% are displayed). Branches in red highlight the three major clades: *S. padanus*, *S. cellostaticus* and *S. scopuliridis*. Leaf labels represent the strain code. The outgroup (*Mycobacterium tuberculosis* H37RV) was manually removed and the branch length information was not incorporated into the tree to facilitate visualization of the bootstrap values.

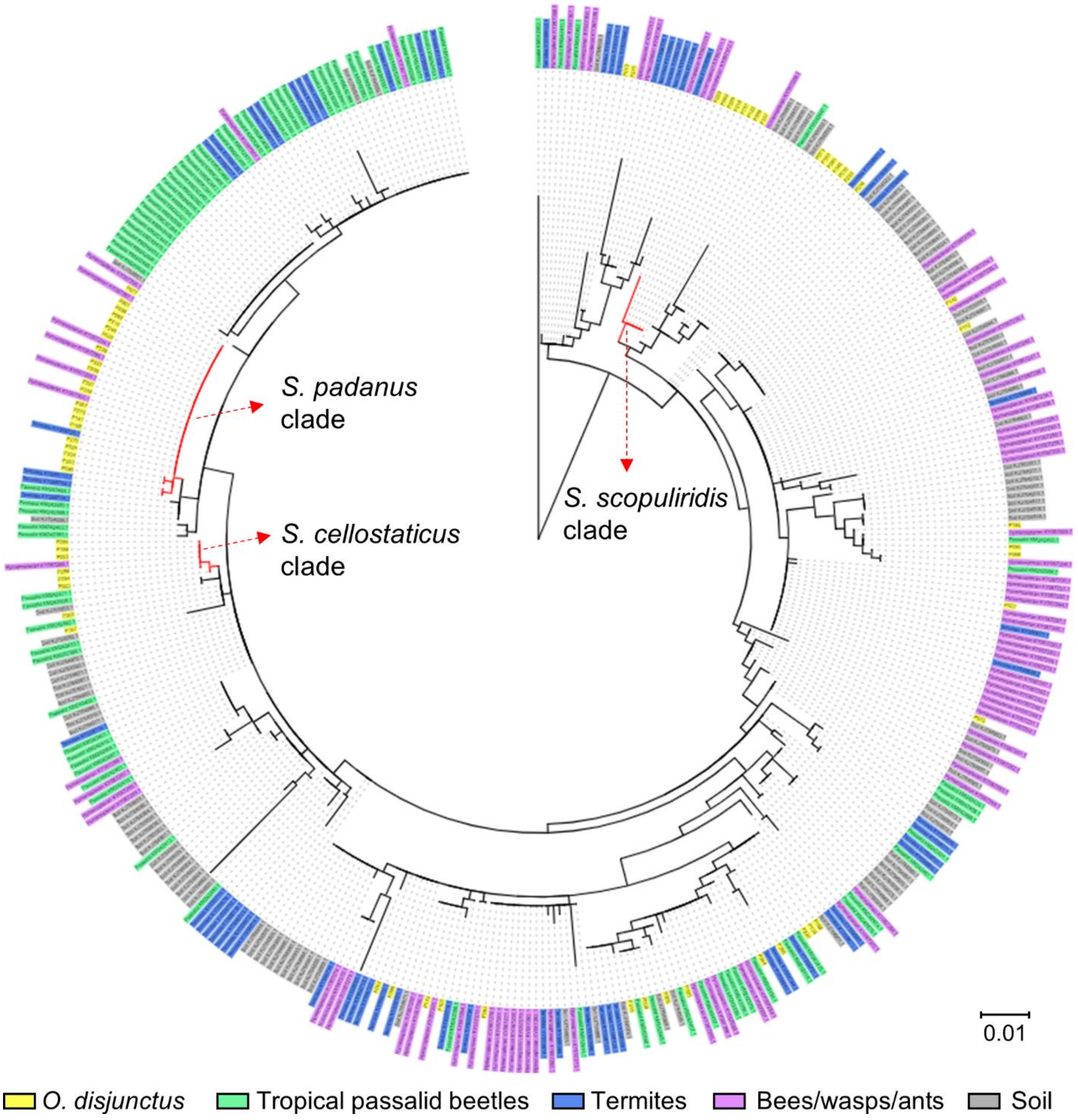


**Figure 4 - figure supplement 8:** Maximum-likelihood phylogenetic tree built using the 16S rRNA gene sequence of *Streptomyces* strains isolated from *O. disjunctus*, tropical passalid beetles, termites, bees/wasps/ants and soil. Scale bar represents branch length in number of substitutions per site. Leaf labels represent the strain code. The outgroup (*Mycobacterium tuberculosis* H37RV) was removed manually from the tree to facilitate visualization. Branches in red highlight the three major clades: *S. padanus*, *S. cellostaticus* and *S. scopuliridis*.

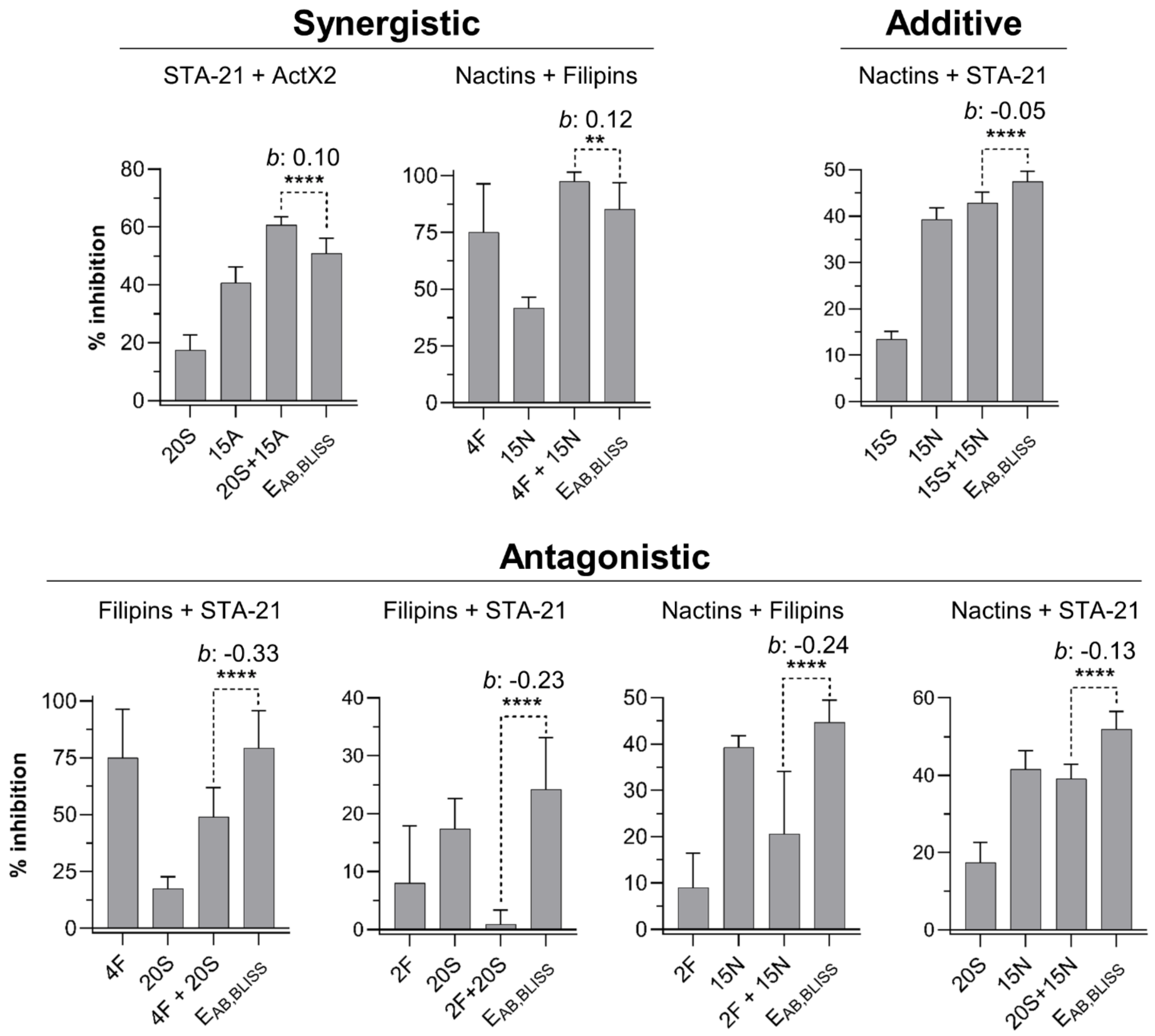


**Figure 5 - figure supplement 1:** Other compound combinations used in the compound interaction assay. Bars represent means (+SD) of percent of growth inhibition (sample size: seven independent biological replicates). Statistical significance was measured using a t-test (****: $p<0.0001$; **: $p=0.009$). Numbers at the X axis represent the tested concentration of each compound in µg/mL (F: filipins. A: actinomycin X2. N: nactins. S: STA-21). *b*: Bliss excess. $E_{AB,BLISS}$: expected value for an independent (additive) interaction between two compounds according to the Bliss Independence model. ActX2: actinomycin X2.

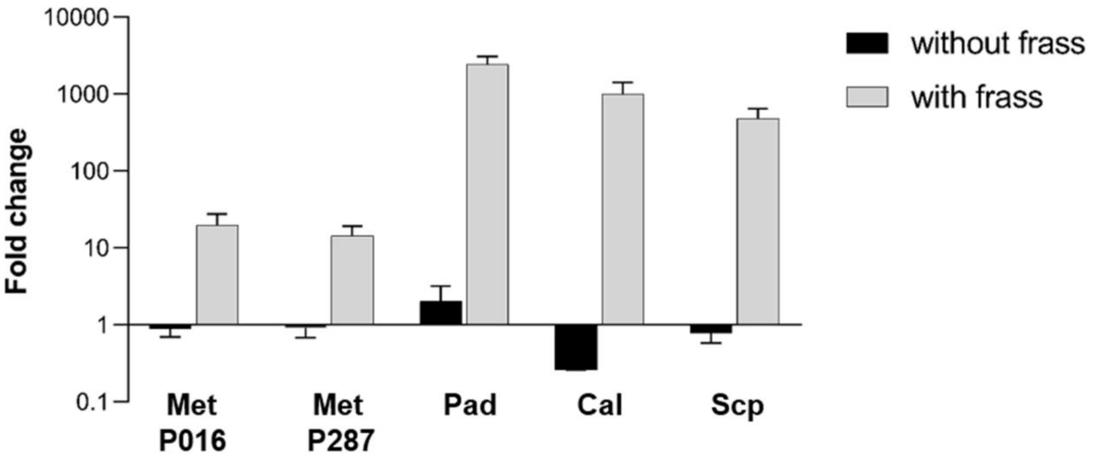


**Figure 6 - figure supplement 1:** All microbes used in the interaction on frass assay were able to use the frass material as a substrate for growth. Bars represent means (+SD) of fold change in growth of each microbe in microtubes with and without frass material after seven days of incubation (compared to the initial inoculum; sample size: eight independent biological replicates). Met P016: *M. anisopliae* P016; Met P287: *M. anisopliae* P287; Pad: *S. padanus* P333; Cal: *S. californicus* P327; Scp: *S. scopuliridis* P239.

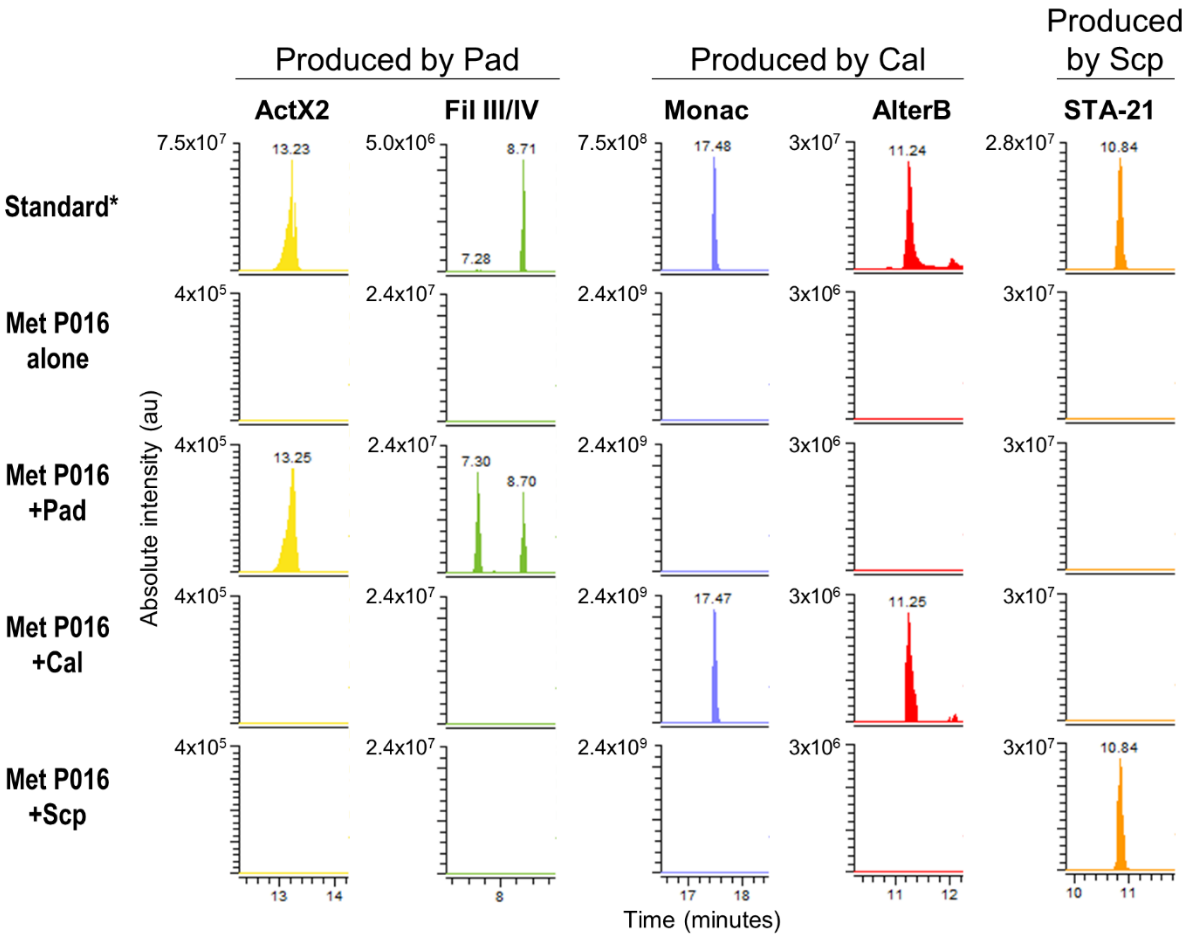


**Figure 6 - figure supplement 2:** EIC of some specialized metabolites detected in treatments containing *M. anisopliae* P016. *Standard: a mixture of crude ethyl acetate extracts of ISP2-solid cultures of the three streptomycetes. Pad: *S. padanus* P333. Cal: *S. californicus* P327. Scp: *S. scopuliridis* P239. Met: *M. anisopliae*. ActX: actinomycin X2. FilIII/IV: filipins III and IV. Monac: monactin. AlterB: alteramide B.

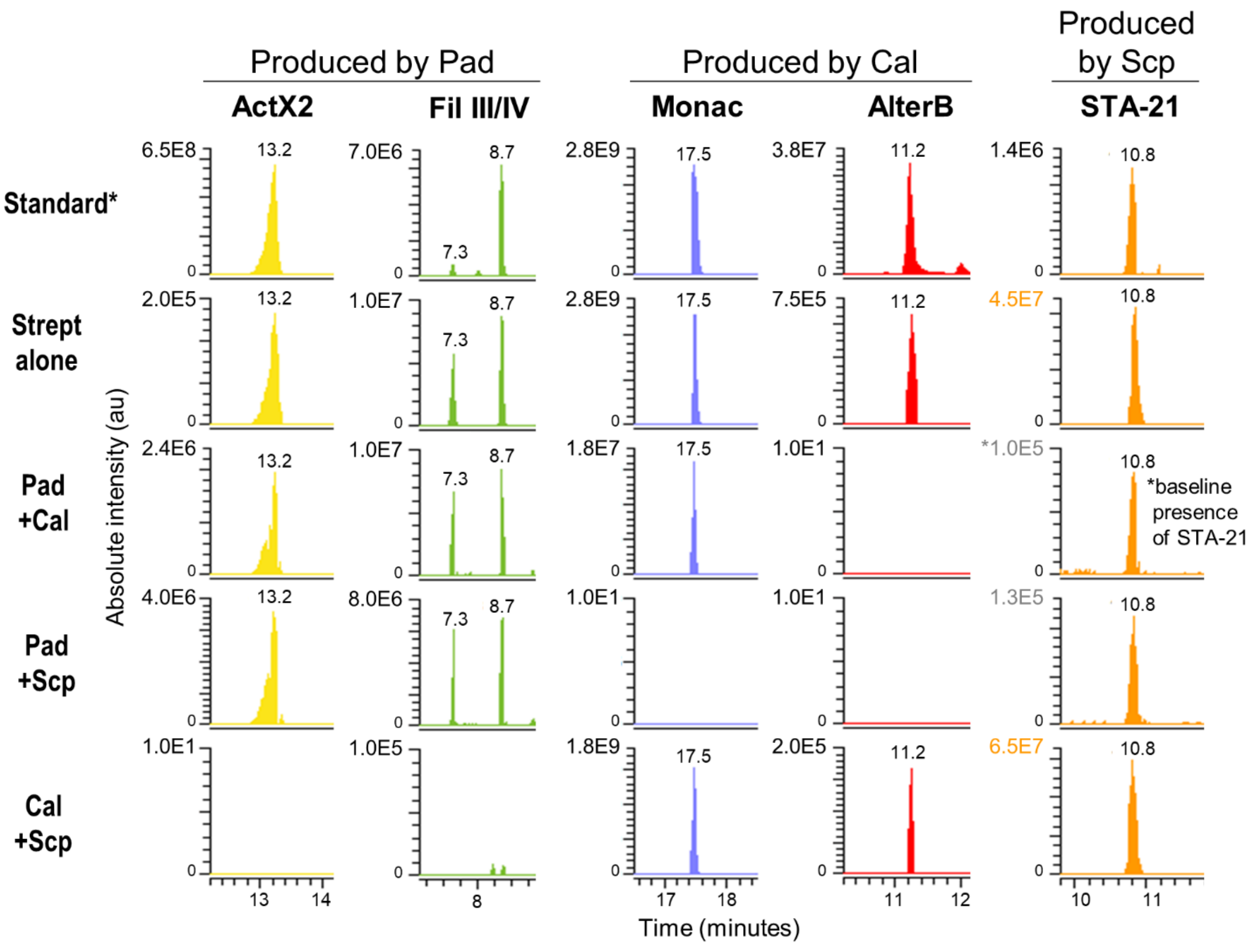


**Figure 6 - figure supplement 3:** EIC of some specialized metabolites detected in treatments containing streptomycetes only. Note that the absolute intensity of the peaks change depending on the combination of microbes. *Standard: a mixture of crude ethyl acetate extracts of ISP2-solid cultures of the three streptomycetes. Pad: *S. padanus* P333. Cal: *S. californicus* P327. Scp: *S. scopuliridis* P239. ActX: actinomycin X2. FilIII/IV: filipins III and IV. Monac: monactin. AlterB: alteramide B. Please note that compound STA-21, produced by Scp, was already present at a low intensity in the frass material since it is an extremely common compound in this environment, therefore, it was detected in all the treatments. We used the intensity observed in the treatment Pad+Cal as its baseline intensity present in the frass prior to the treatments. Note that its intensity in the treatment Pad+Scp is very close to the baseline, whereas its intensity in the treatment Cal+Scp is similar to the one observed for Scp alone.
