## Appendix Figures for "Multiple lineages of *Streptomyces* produce antimicrobials within passalid beetle galleries across eastern North America"

**(1) Actinomycin D**

$[M+H]^+$

Mass error: 1.7 ppm

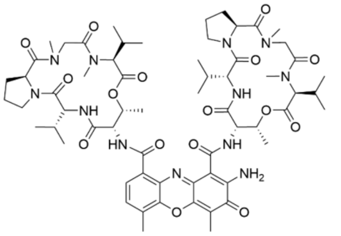

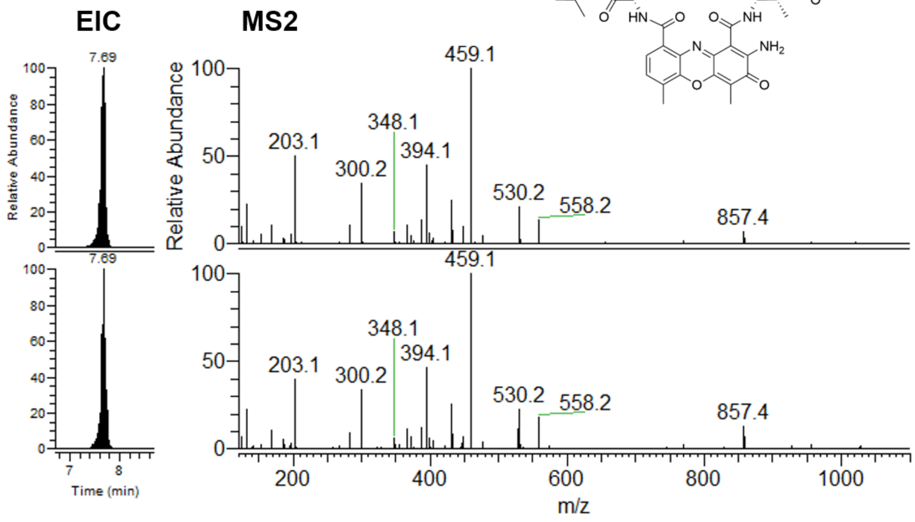


**Standard**

NL: 1.36E6
021620_Std_ActD#3128-3223 RT: 7.63-7.72 AV: 2 F: FTMS + p ESI d Full ms2 1255.6366@hcd30.00 [50.0000-1300.0000]

**Strain P333**

NL: 6.63E6
011319_EtAcMS2_P333#3913-4025 RT: 7.67-7.77 AV: 2 F: FTMS + p ESI d Full ms2 1255.6366@hcd30.00 [50.0000-1300.0000]

**(2) Actinomycin X2**

$[M+H]^+$

Mass error: 2.2 ppm

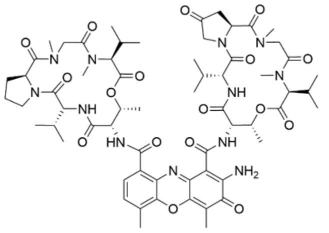

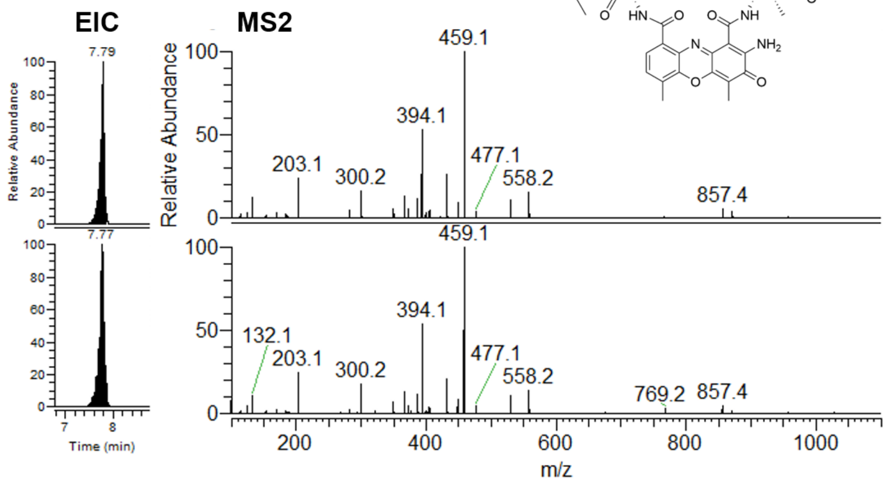


**Standard**

NL: 1.61E6
052120_Std_ActinomycinX2#3188-3253 RT: 7.74-7.82 AV: 2 F: FTMS + p ESI d Full ms2 1269.6119@hcd30.00 [50.0000-1315.0000]

**Strain P333**

NL: 1.90E7
011319_EtAcMS2_P333#3969-4052 RT: 7.74-7.84 AV: 2 F: FTMS + p ESI d Full ms2 1269.6119@hcd30.00 [50.0000-1315.0000]

**(3) STA-21**

[M+H]$^+$

Mass error: 1.6 ppm

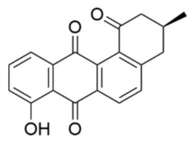


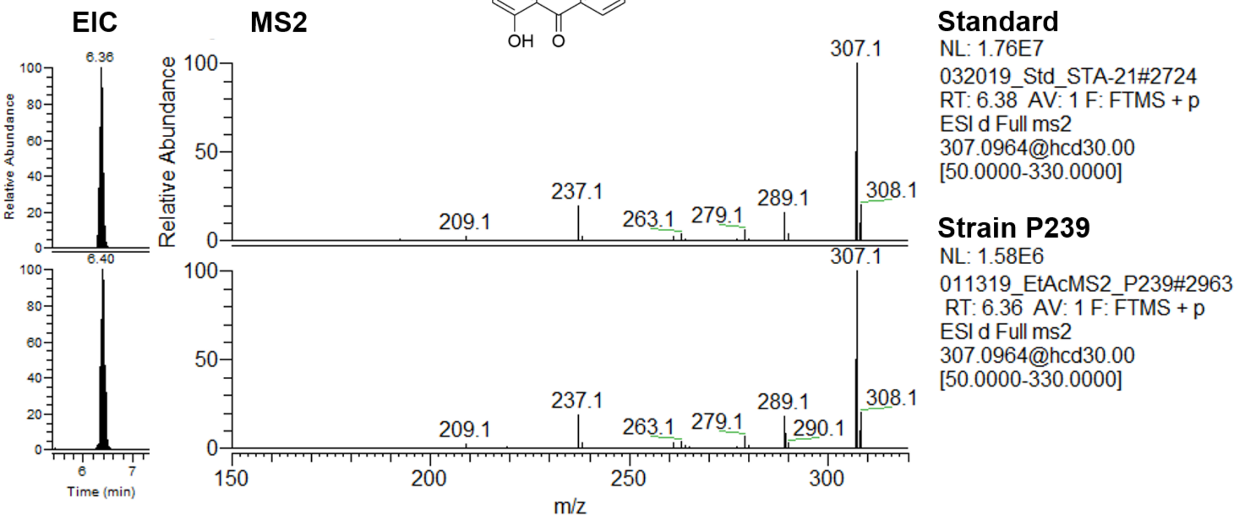


**(4) Rubiginone B2**

[M+H]$^+$

Mass error: 2.2 ppm

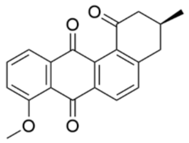

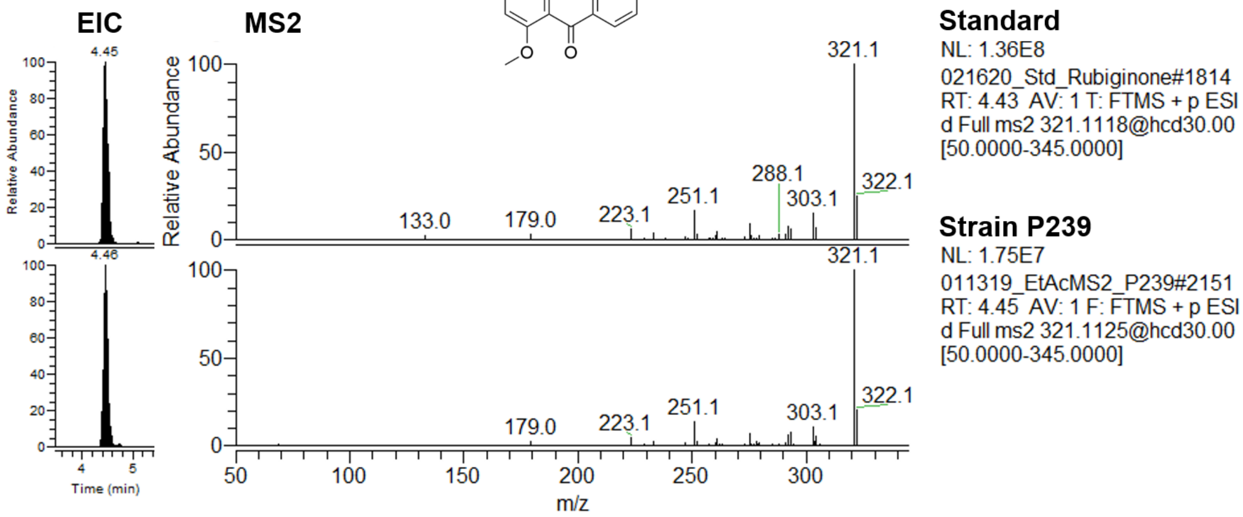

**(5) Cycloheximide**

$[M+H]^+$

Mass error: 2.1 ppm

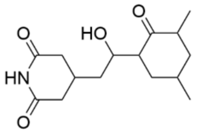


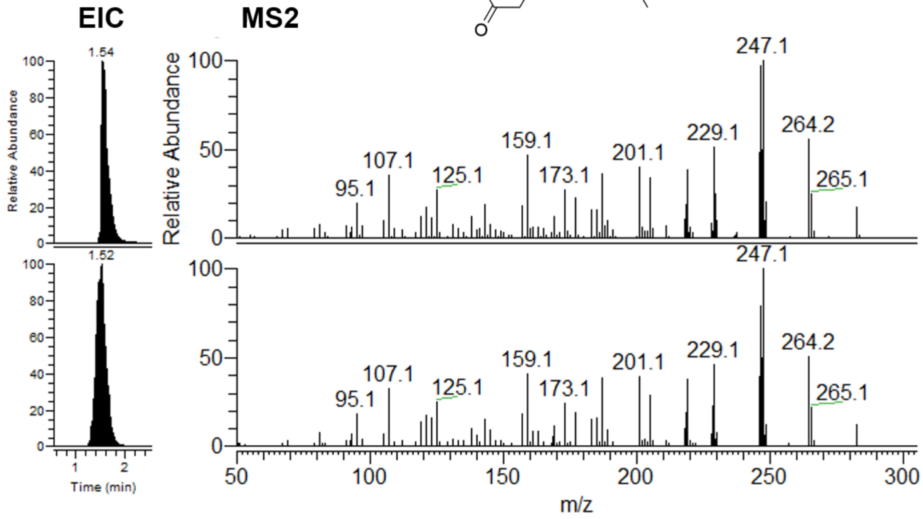


**Standard**

NL: 1.34E8

012419_cycloheximide-#802

RT: 1.62 AV: 1 F: FTMS + p

ESI d Full ms2

282.1703@hcd30.00

[50.0000-305.0000]

**Strain P263**

NL: 3.05E7

011319_EtAcMS2_P263#816

RT: 1.58 AV: 1 F: FTMS + p

ESI d Full ms2

282.1703@hcd30.00

[50.0000-305.0000]

**(6) Nonactin**

$[M+Na]^+$

Mass error: 2.6 ppm

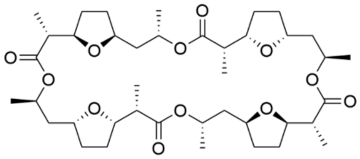

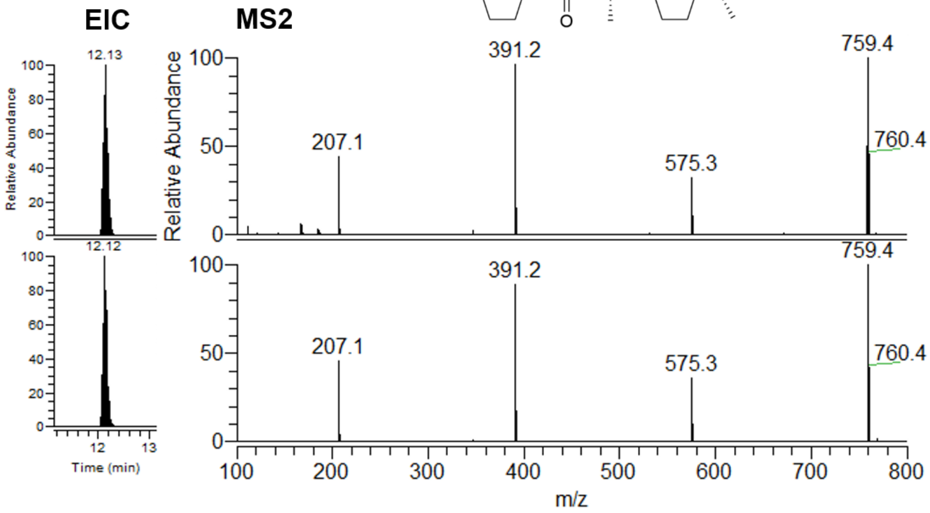


**Standard**

NL: 1.18E7

092520_Nactins_Std#5285 RT: 12.08 AV: 1 F: FTMS + p ESI d Full ms2 759.4280@hcd30.00 [50.0000-795.0000]

**Strain P327**

NL: 4.89E6

011319_EtAcMS2_P327#5713

RT: 12.08 AV: 1 F: FTMS + p

ESI d Full ms2

759.4280@hcd30.00

[50.0000-795.0000]

**(7) Monactin**

$[M+Na]^+$

Mass error: 3.2 ppm

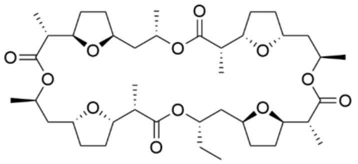

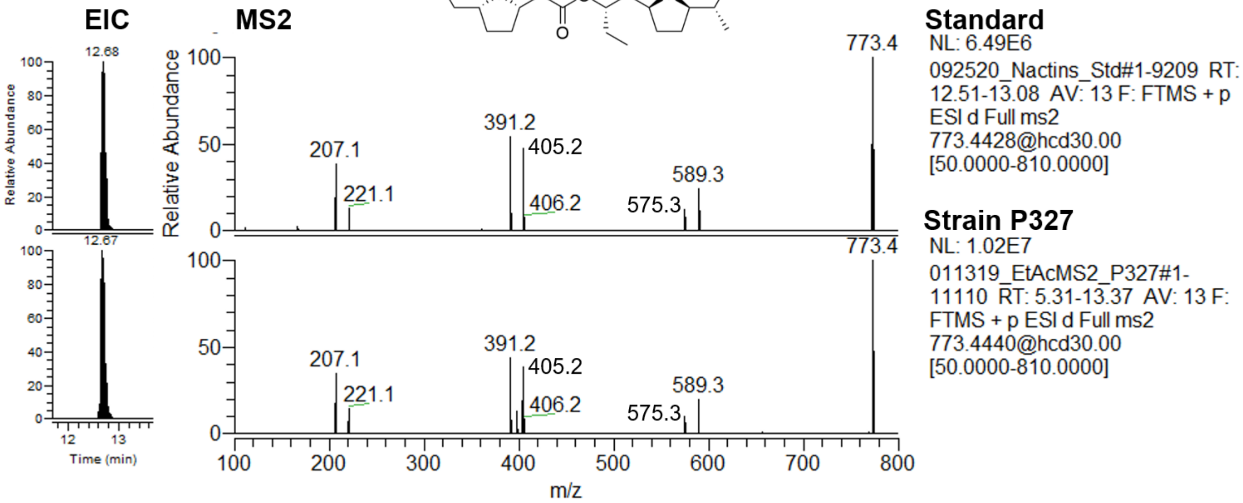


**(8) Dinactin**

$[M+Na]^+$

Mass error: 3.4 ppm

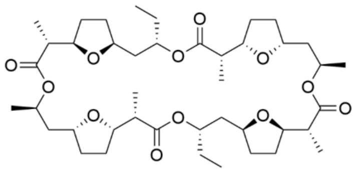

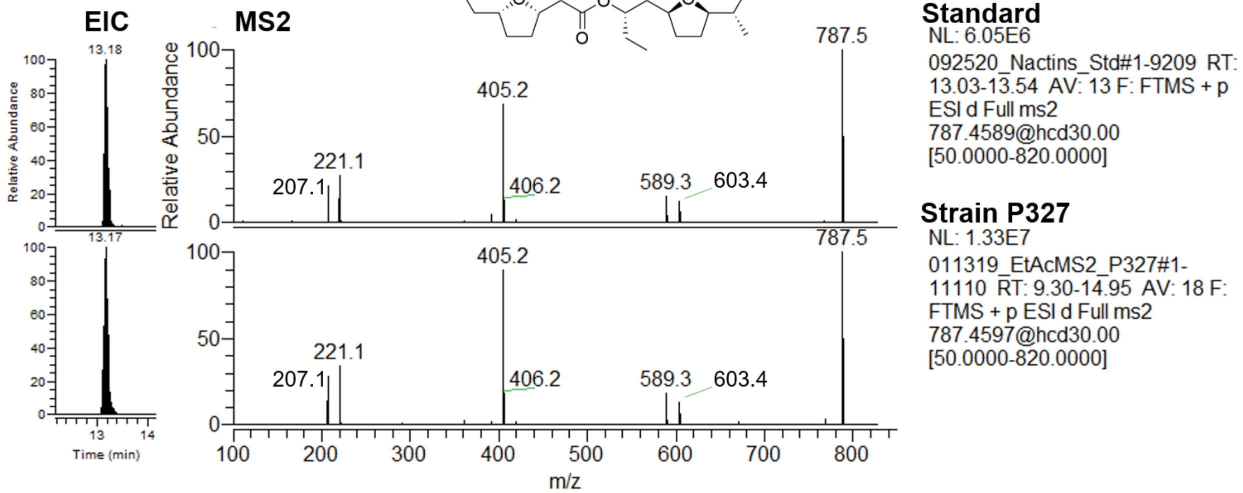

**(9) Trinactin**

$[M+Na]^+$

Mass error: 2.5 ppm

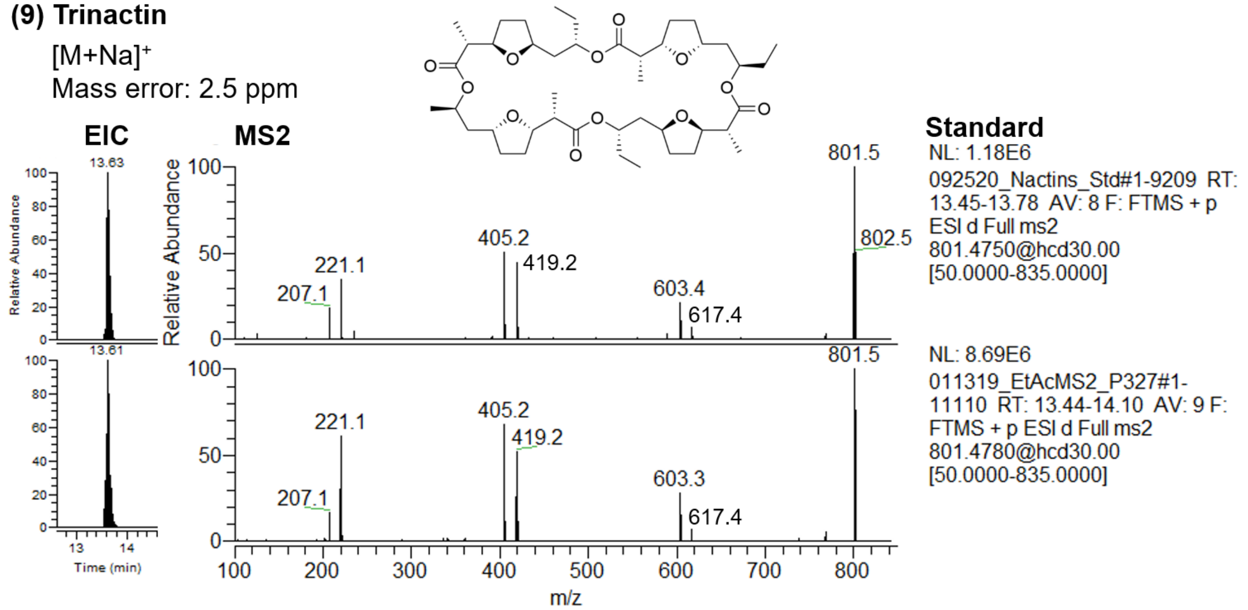


**(10) Tetranactin**

$[M+Na]^+$

Mass error: 1.7 ppm

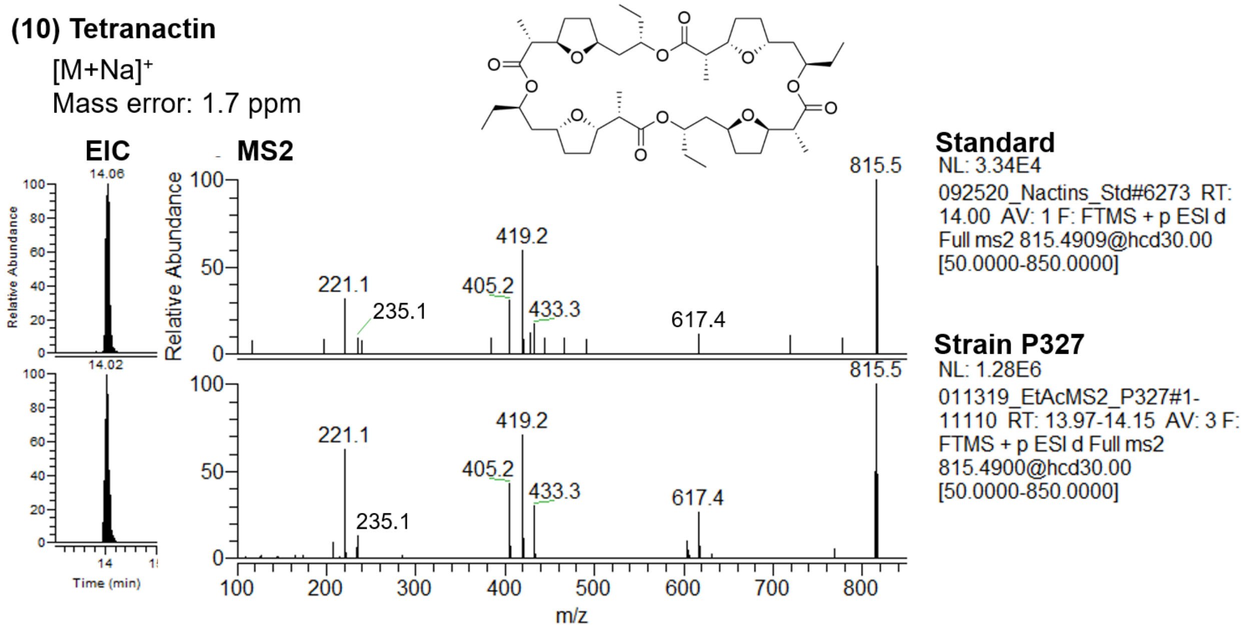

**(11) Filipin I**

$[M+Na]^+$

Mass error: 1.1 ppm

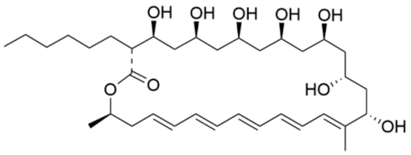


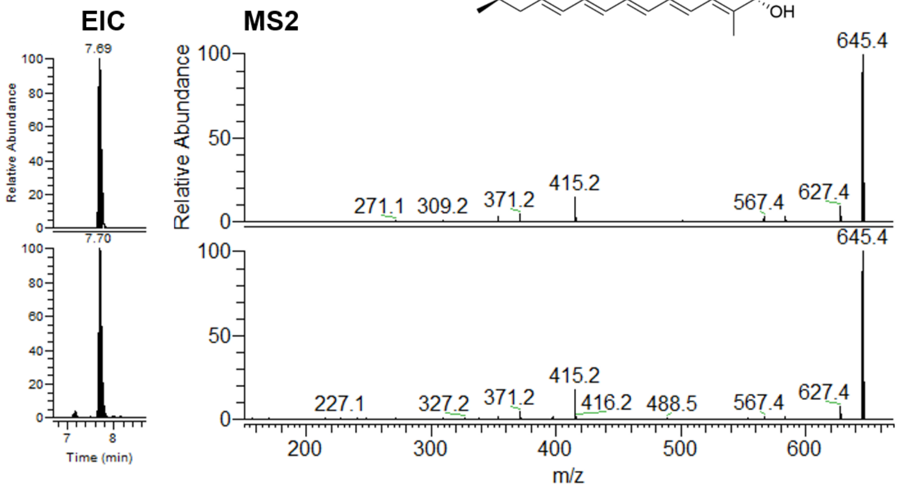


**Standard**

NL: 4.28E5

052120_Std_FilipinComplex#3604-3691 RT: 7.64-7.73 AV: 2 F: FTMS + p ESI d Full ms2 645.3947@hcd30.00 [50.0000-675.0000]

**Strain P181**

NL: 1.29E5

021320_EtAc_P181#3522-3606 RT: 7.65-7.74 AV: 2 F: FTMS + p ESI d Full ms2 645.3947@hcd30.00 [50.0000-675.0000]

**(12) Filipin II**

$[M+Na]^+$

Mass error: 1.7 ppm

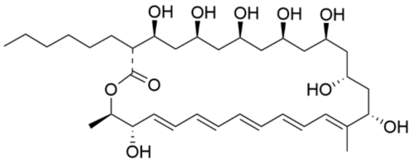


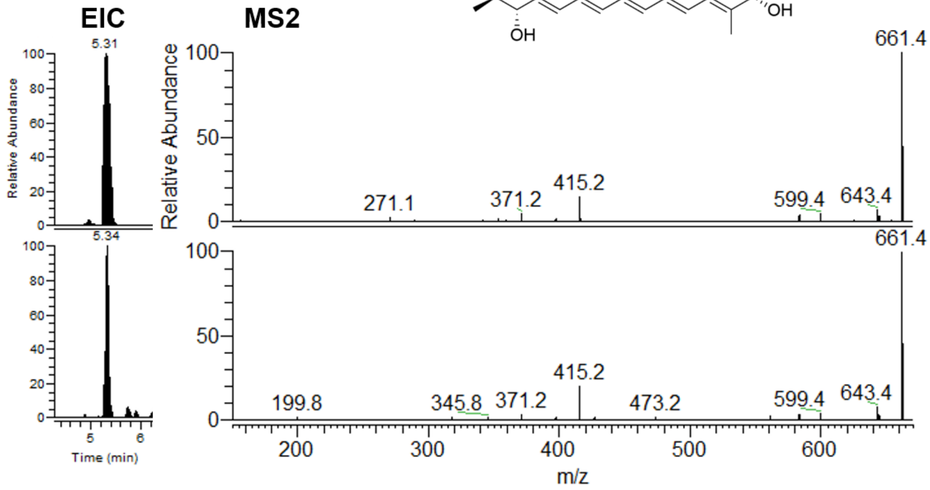


**Standard**

NL: 5.50E6

052120_Std_FilipinComplex#2590 RT: 5.32 AV: 1 F: FTMS + p ESI d Full ms2 661.3928@hcd30.00 [50.0000-695.0000]

**Strain P181**

NL: 1.30E5

021320_EtAc_P181#2491 RT: 5.36 AV: 1 F: FTMS + p ESI d Full ms2 661.3928@hcd30.00 [50.0000-695.0000]

**(13) Filipin III**

$[M+Na]^+$

Mass error: 0.3 ppm

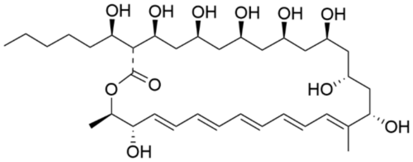


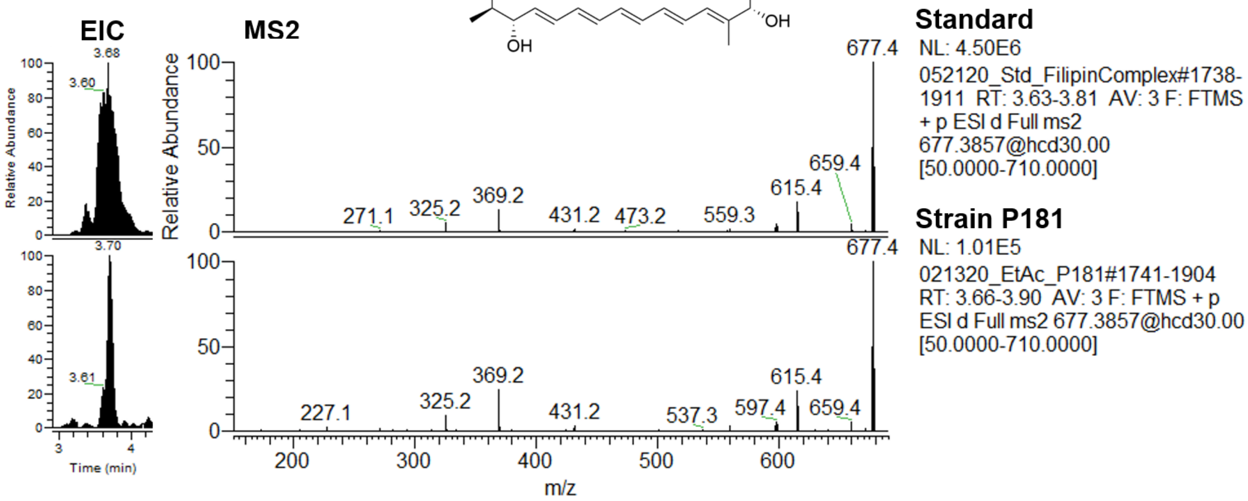


**(18) Antimycin A**

$[M+H]^+$

Mass error: 0.0 ppm

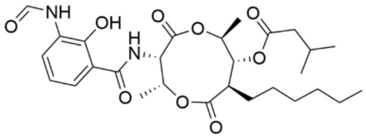


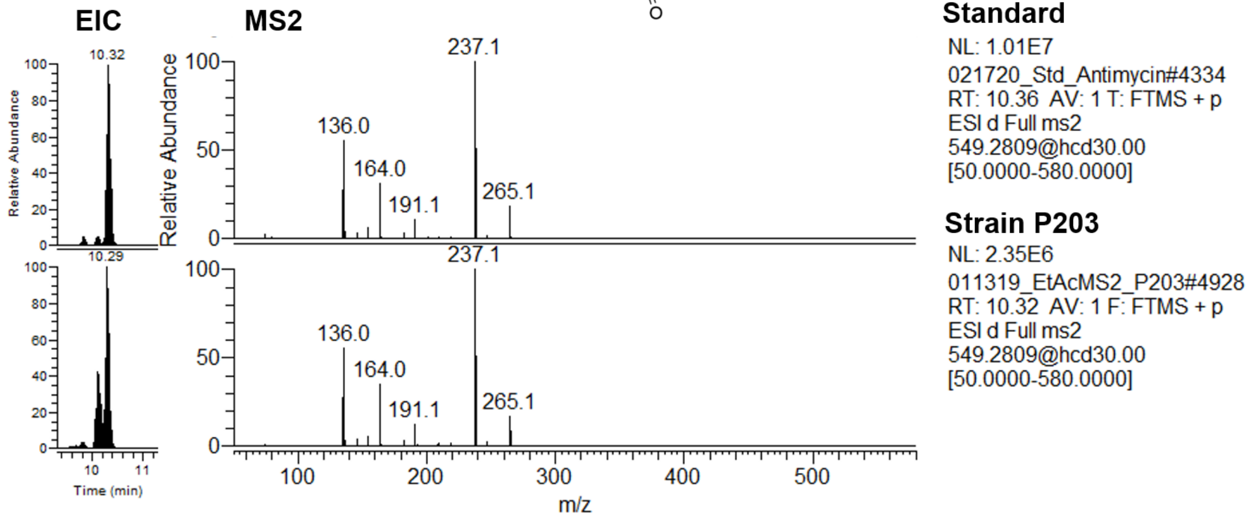

**(19) Nocardamine**

$[M+H]^+$

Mass error: 0.8 ppm

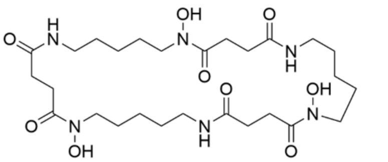

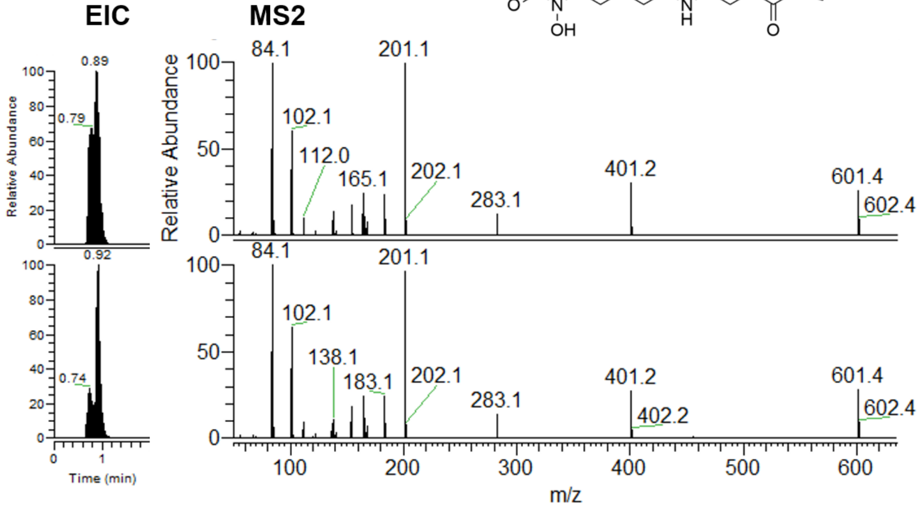


**Standard**

NL: 9.53E5

022520_Std_Nocardamine#207-442 RT: 0.70-0.96 AV: 4 F: FTMS + p ESI d Full ms2 601.3552@hcd30.00 [50.0000-630.0000]

**Strain P307**

NL: 9.62E5

011319_EtAcMS2_P307#207-465 RT: 0.70-0.98 AV: 4 F: FTMS + p ESI d Full ms2 601.3552@hcd30.00 [50.0000-630.0000]

**(20) Bafilomycin A1**

$[M+Na]^+$

Mass error: 1.7 ppm

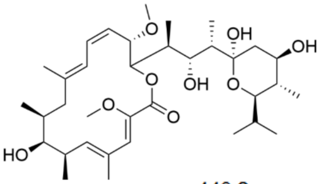

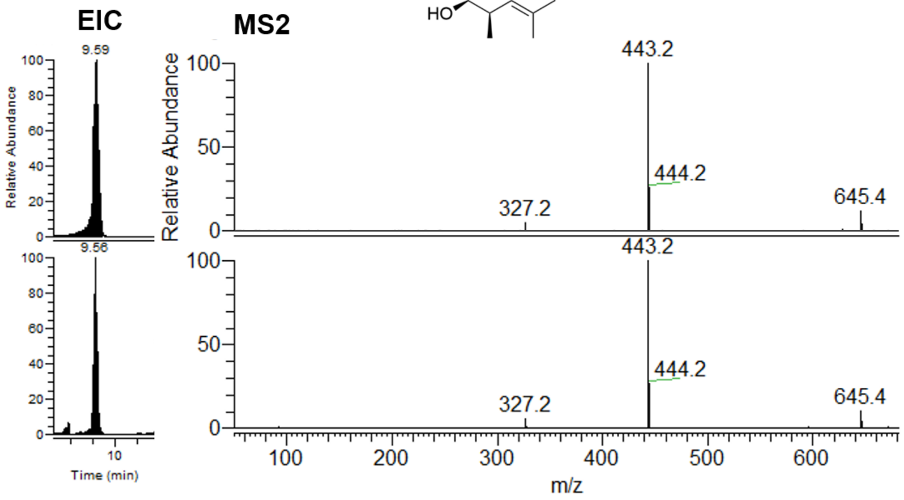


**Standard**

NL: 6.33E6

100219_BafilomycinA1#3952-4066 RT: 9.54-9.63 AV: 2 F: FTMS + c ESI d Full ms2 645.3981@hcd30.00 [50.0000-675.0000]

**Strain P059**

NL: 5.72E5

021320_EtAc_P059#4397-4507 RT: 9.51-9.60 AV: 2 F: FTMS + p ESI d Full ms2 645.3964@hcd30.00 [50.0000-675.0000]

**(21) Bafilomycin B1**

[M+Na]+

Mass error: 2.0 ppm

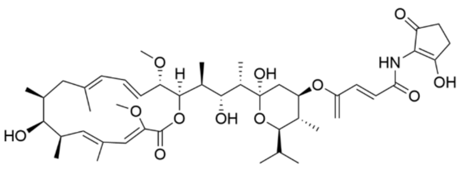

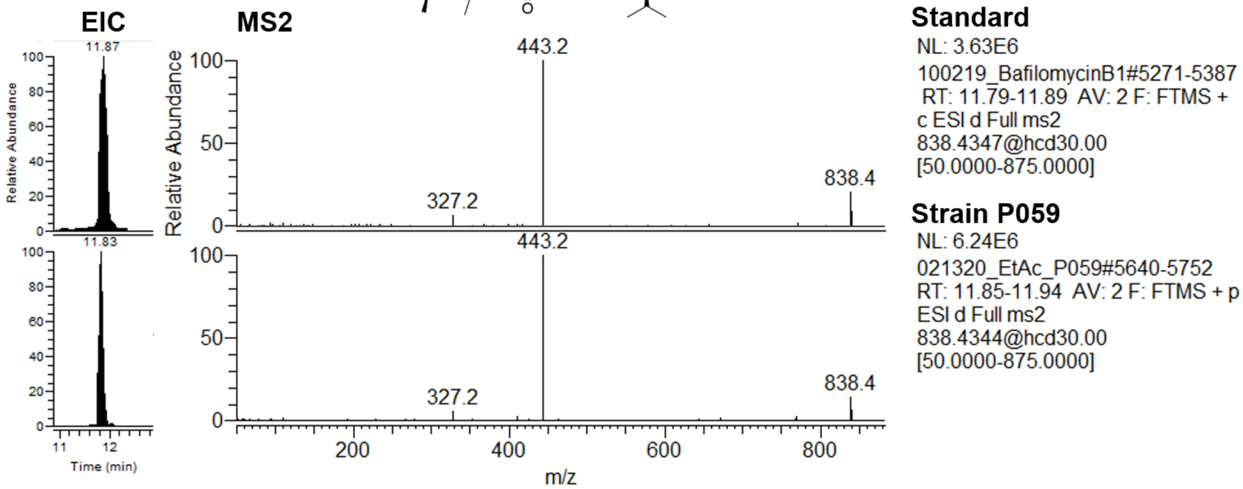


**(22) Novobiocin**

[M+H]+

Mass error: 2.4 ppm

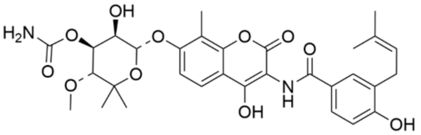

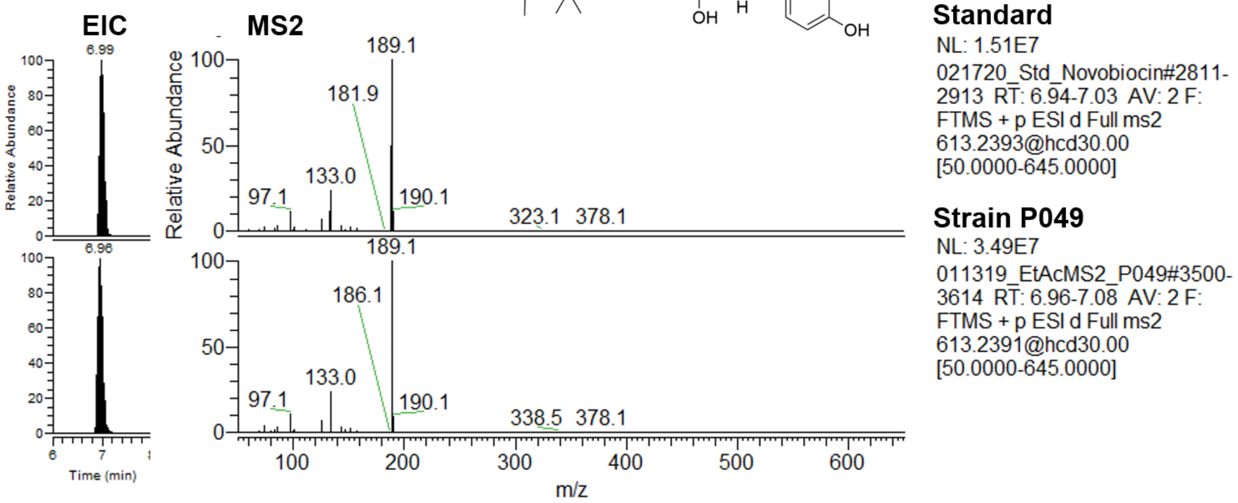

**(24) Piericidin A**

$[M+H]^+$

Mass error: 2.6 ppm

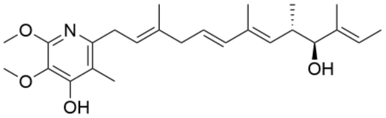


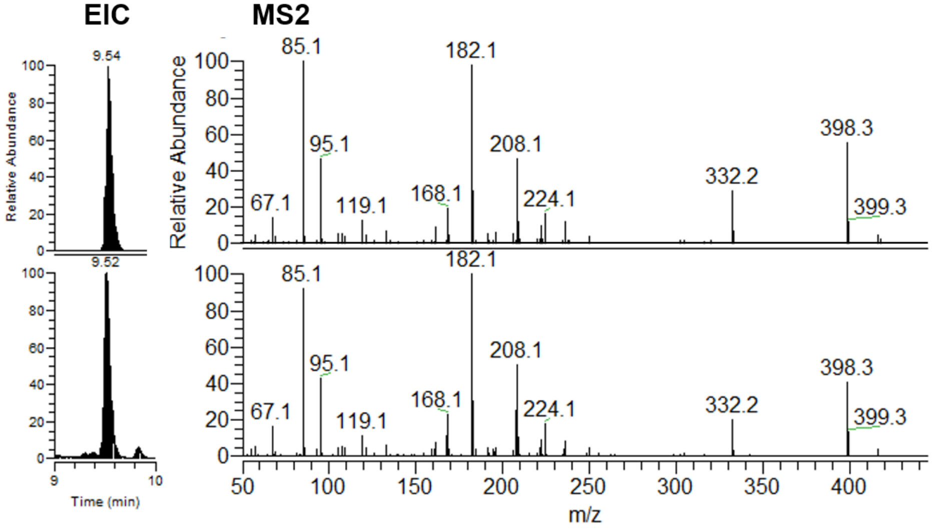


**Standard**

NL: 2.04E7

100219_Piericidin#4783 RT: 9.53 AV: 1 F: FTMS + c ESI d Full ms2 416.2798@hcd30.00 [50.0000-445.0000]

**Strain P190**

NL: 2.29E6

021420_EtAc_P190#4166 RT: 9.51 AV: 1 T: FTMS + p ESI d Full ms2 416.2788@hcd30.00 [50.0000-445.0000]

**(25) Nigericin**

$[M+Na]^+$

Mass error: 0.5 ppm

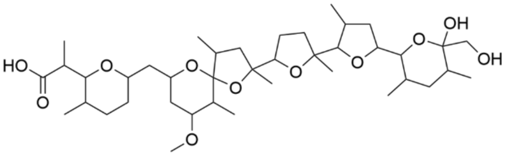


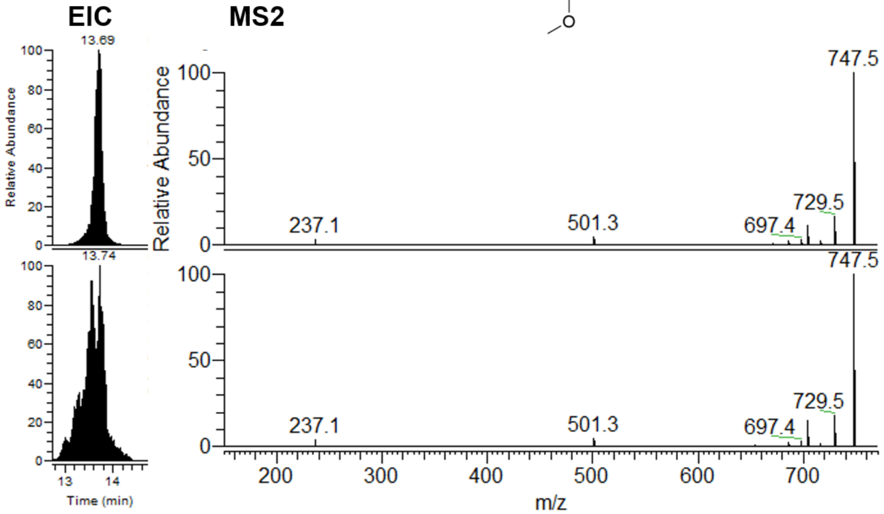


**Standard**

NL: 1.28E8

052120_Std_Nigericin#5740-5841 RT: 13.64-13.72 AV: 2 F: FTMS + p ESI d Full ms2 747.4639@hcd30.00 [50.0000-780.0000]

**Strain P009**

NL: 1.98E8

011319_EtAcMS2_P009#7502-7611 RT: 13.61-13.79 AV: 3 F: FTMS + p ESI d Full ms2 747.4639@hcd30.00 [50.0000-780.0000]

**(26) Beauvericin**
[M+Na]+
Mass error: 0.2 ppm

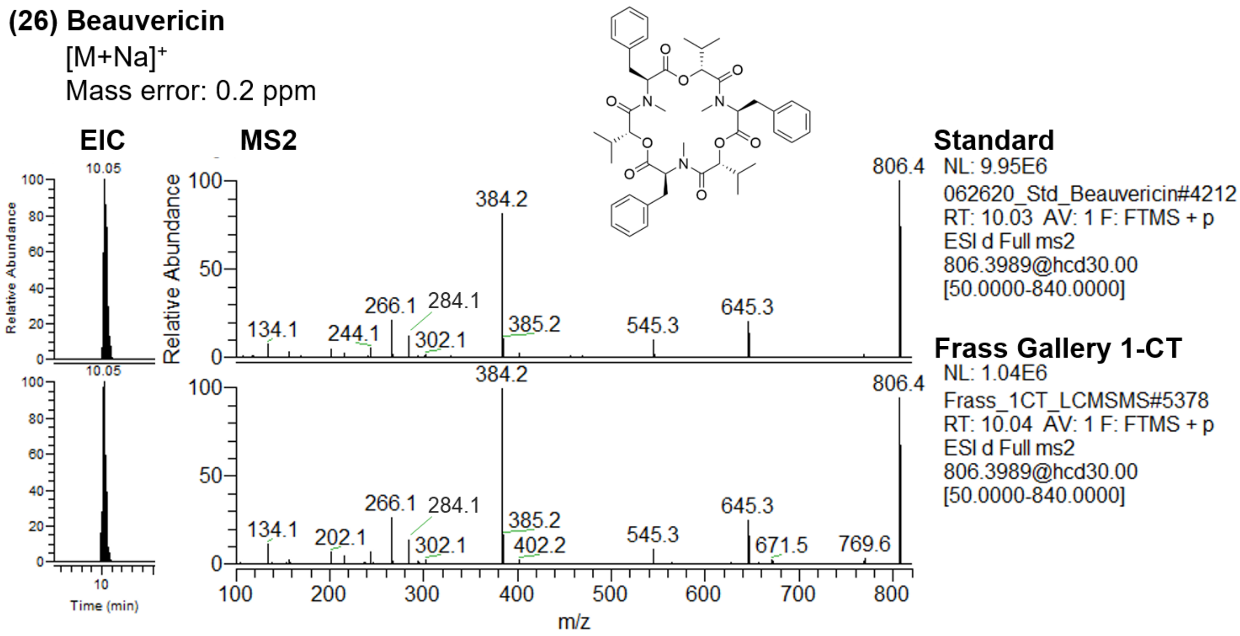


**Appendix 1 – Figure 1:** Extracted ion chromatogram (EIC, showing retention time) and MS2 spectra of each compound annotated at identification level 1 (Actinomycin D, Actinomycin X2, STA-21, Rubiginone B2, Cycloheximide, Nonactin, Monactin, Dinactin, Trinactin, Tetranactin, Filipin I, Filipin II, Filipin III, Antimycin A, Nocardamine, Bafilomycin A1, Bafilomycin B1, Novobiocin, Piericidin A, Nigericin, Beauvericin), comparing a commercial standard (top) to the culture extract of an exemplary microbe or environmental frass extract (bottom).

**(16) Alteramide A**

$[M+H]^+$

Mass error: 2.2 ppm

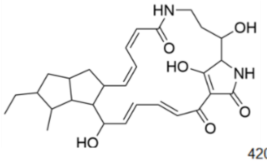


**Strain P327, MS2 spectrum of *m/z* 511.2786**

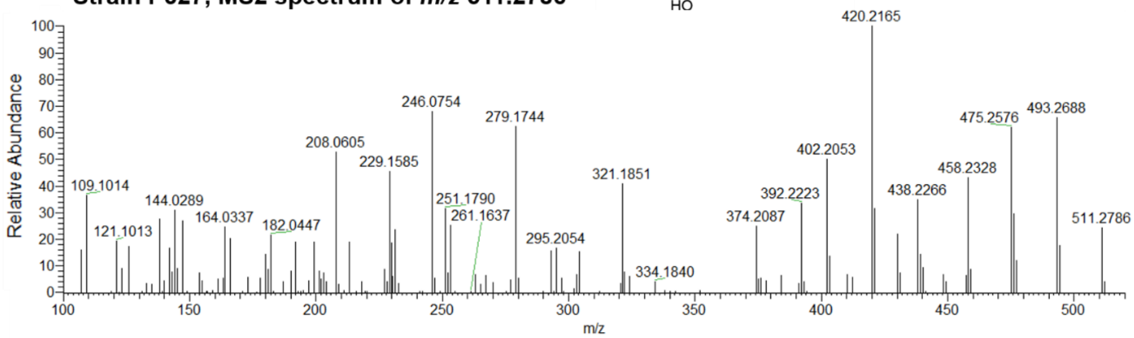


***Streptomyces albus* J1074, MS2 spectrum of *m/z* 511.2800**

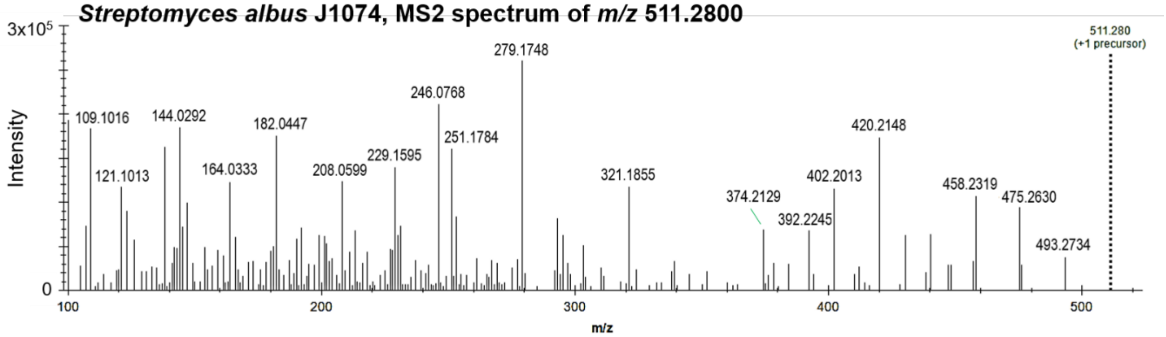


**(17) Alteramide B**

$[M+H]^+$

Mass error: 4.0 ppm

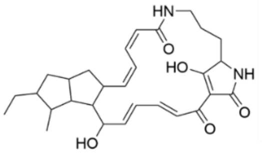


**Strain P327, MS2 spectrum of *m/z* 495.2856**

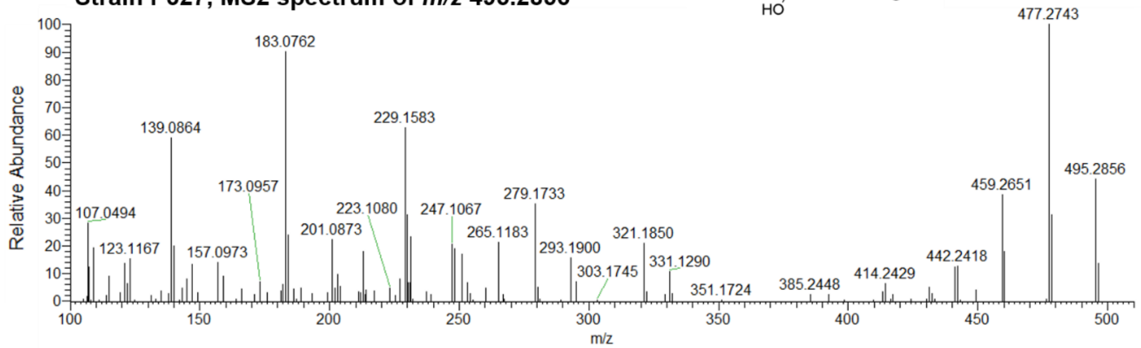


***Streptomyces albus* J1074, MS2 spectrum of *m/z* 495.2848**

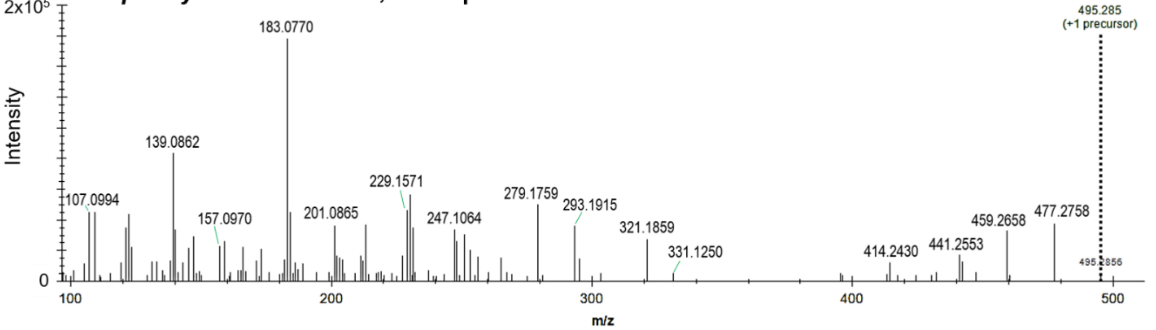

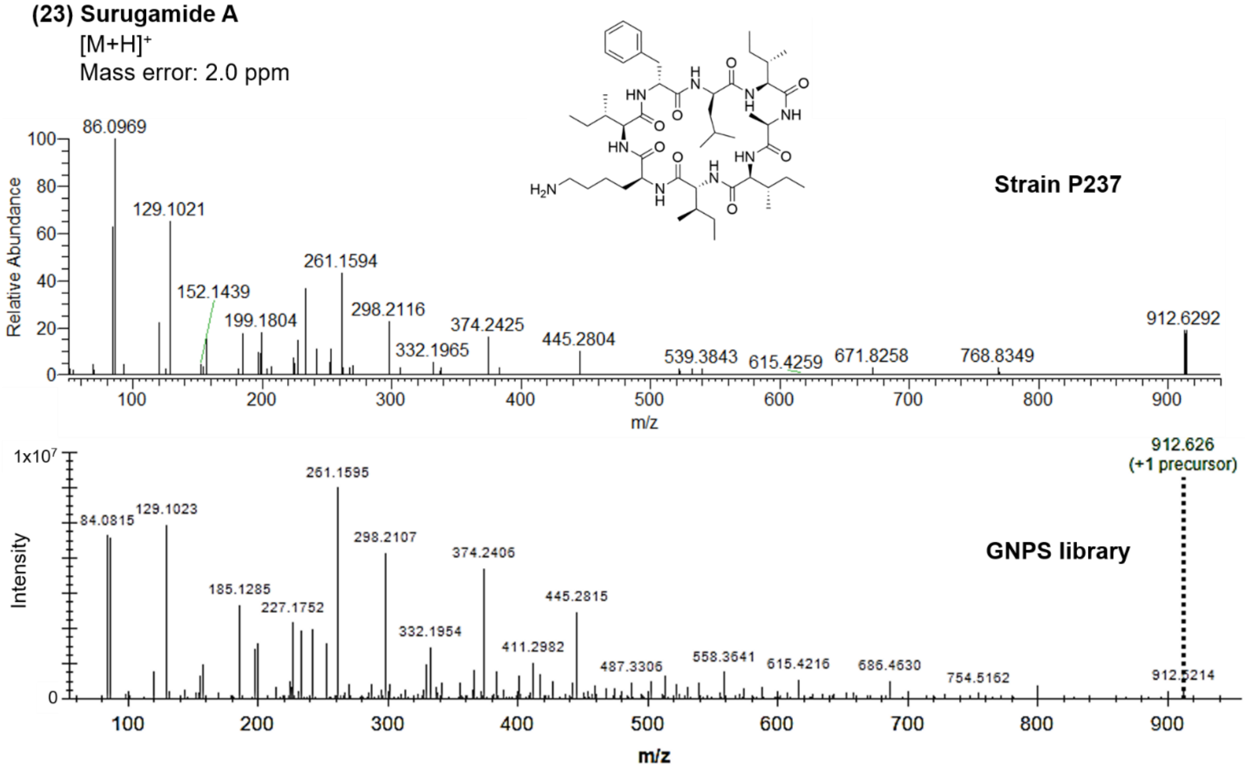


**Appendix 1 - Figure 2:** MS2 spectra of each compound annotated at identification level 2 (Alteramide A, Alteramide B, Surugamide A), comparing a spectrum detected in the culture extract of an exemplary microbe (top) to a publicly available spectrum on the MassIVE repository (bottom). Publicly available spectra can be found at: Alteramide A, Alteramide B: f.MSV000079516/ccms_peak/Labelled/R5_lab_J1074_pre.mzXML; Surugamide A: f.MSV000079519/ccms_peak/Unlabelled/A1_unlab_J1074_pre.mzXML (accessed on June/2020).

**(14) Filipin IV**

[M+Na]+

Mass error: 1.0 ppm

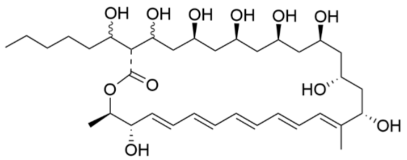


**MS2**

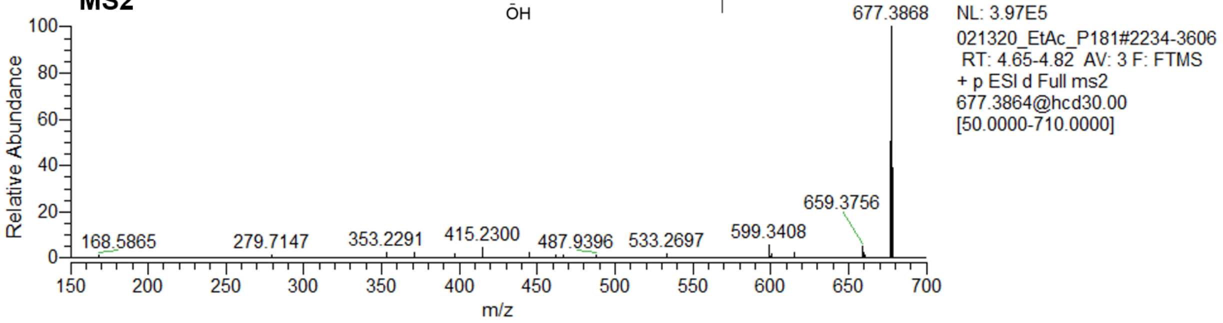


**(15) Fungichromin**

[M+Na]+

Mass error: 2.2 ppm

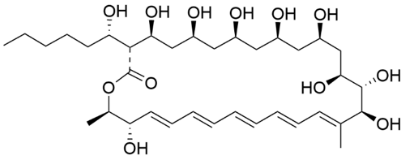


**MS2**

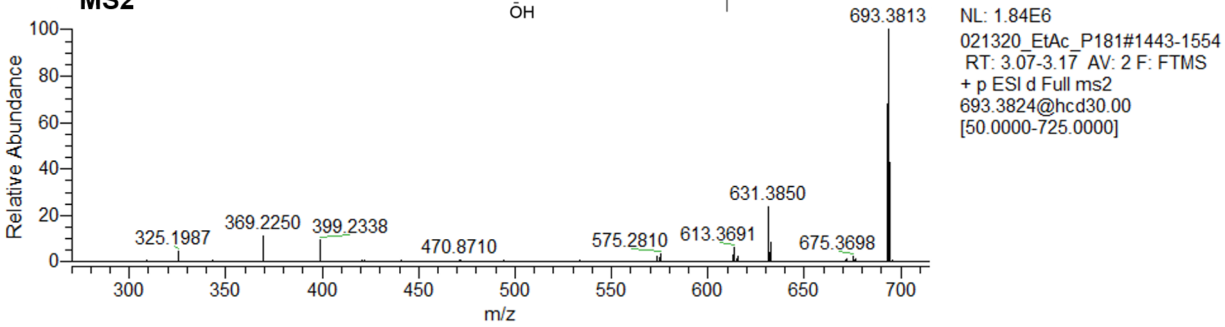


**Appendix 1 - Figure 3:** MS2 spectrum of each compound putatively annotated at identification level 3 (Filipin IV, Fungichromin), detected in the culture extract of an exemplary microbe. Annotations were made based on fragmentation similarities with other analogs of the same family annotated at identification level 1 (filipins I-III).

**A) Pupal chamber material (gallery 17-LA)**

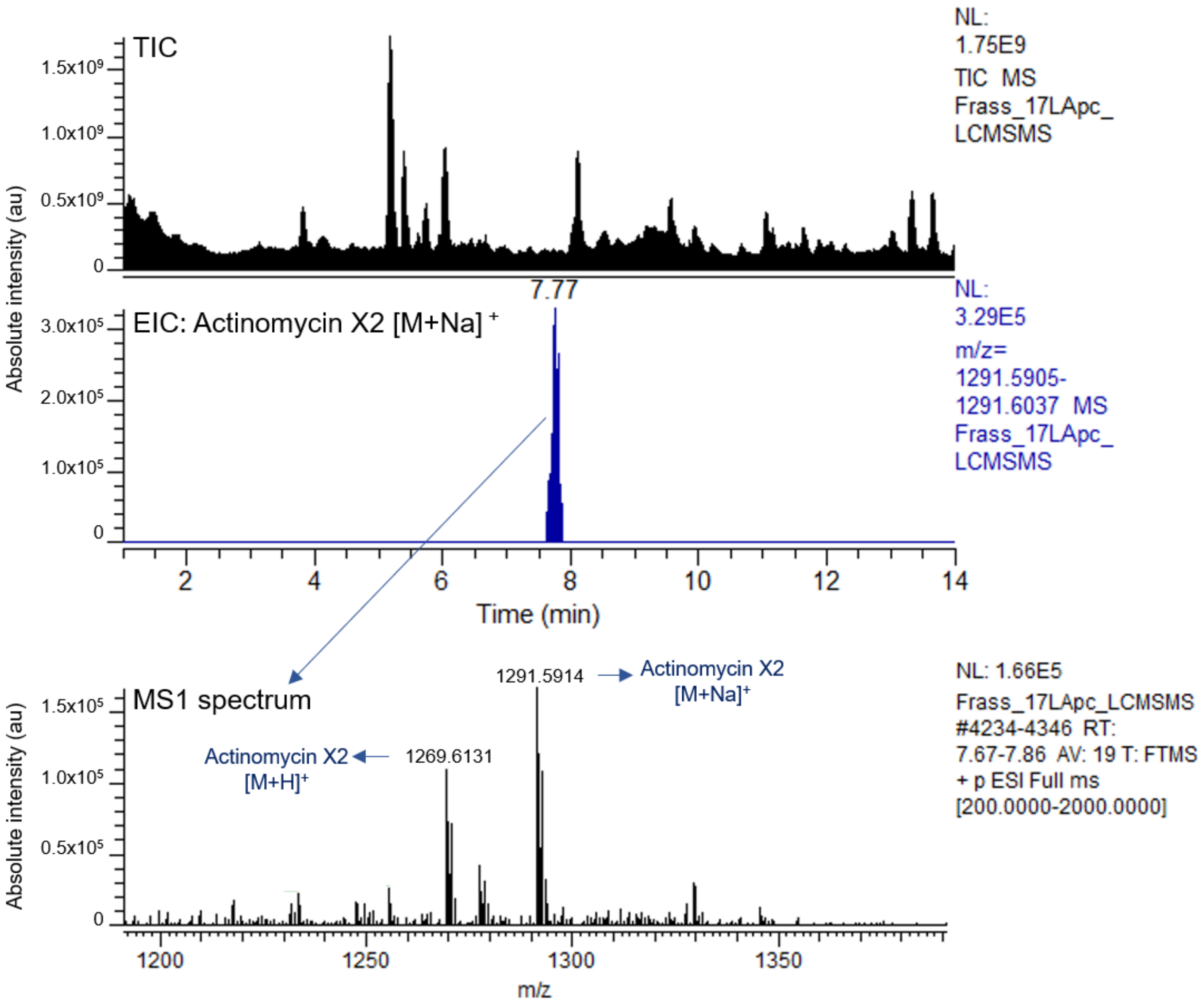

**B) Old frass (gallery 3-DC)**

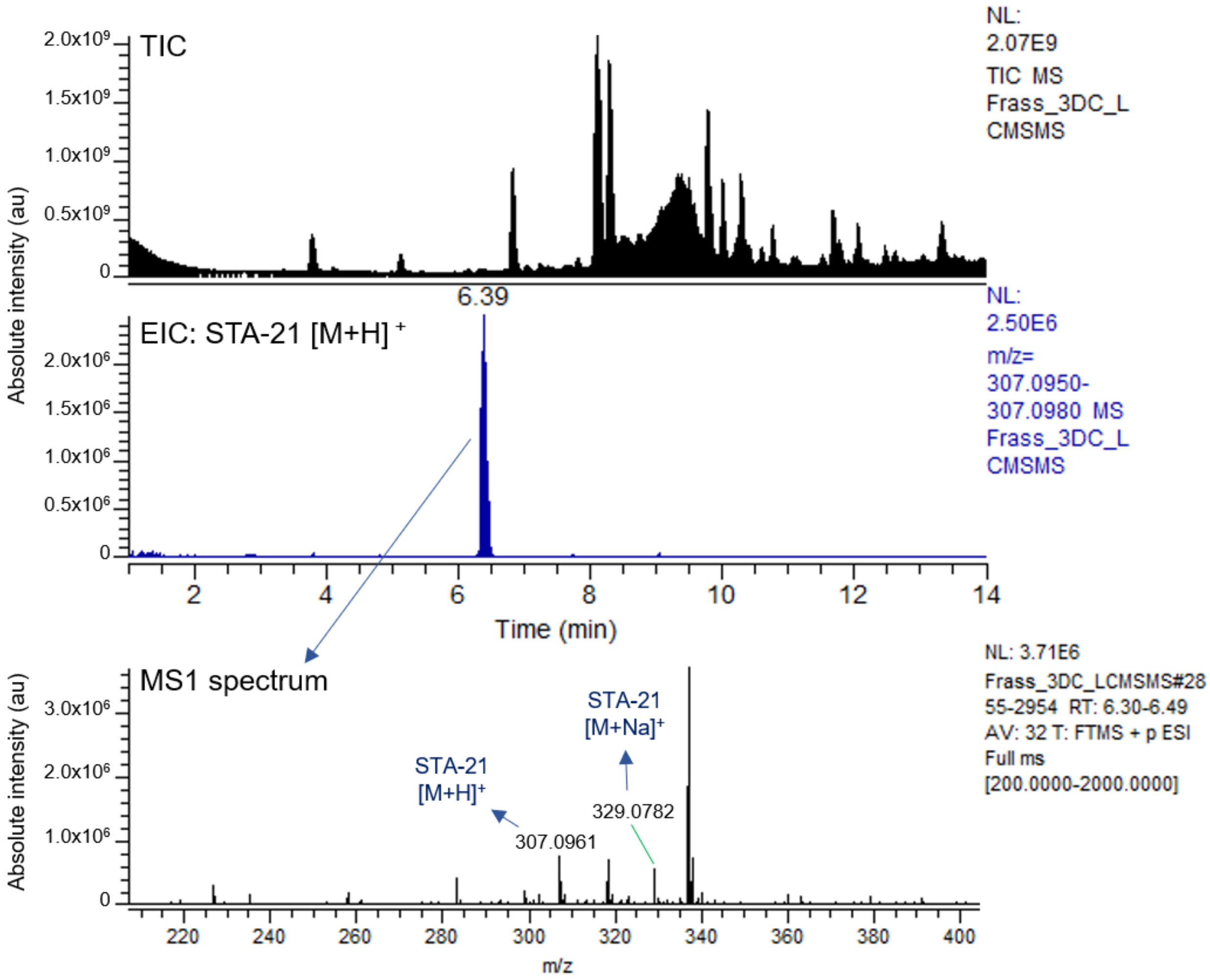

**C) Old frass (gallery 20-LA)**

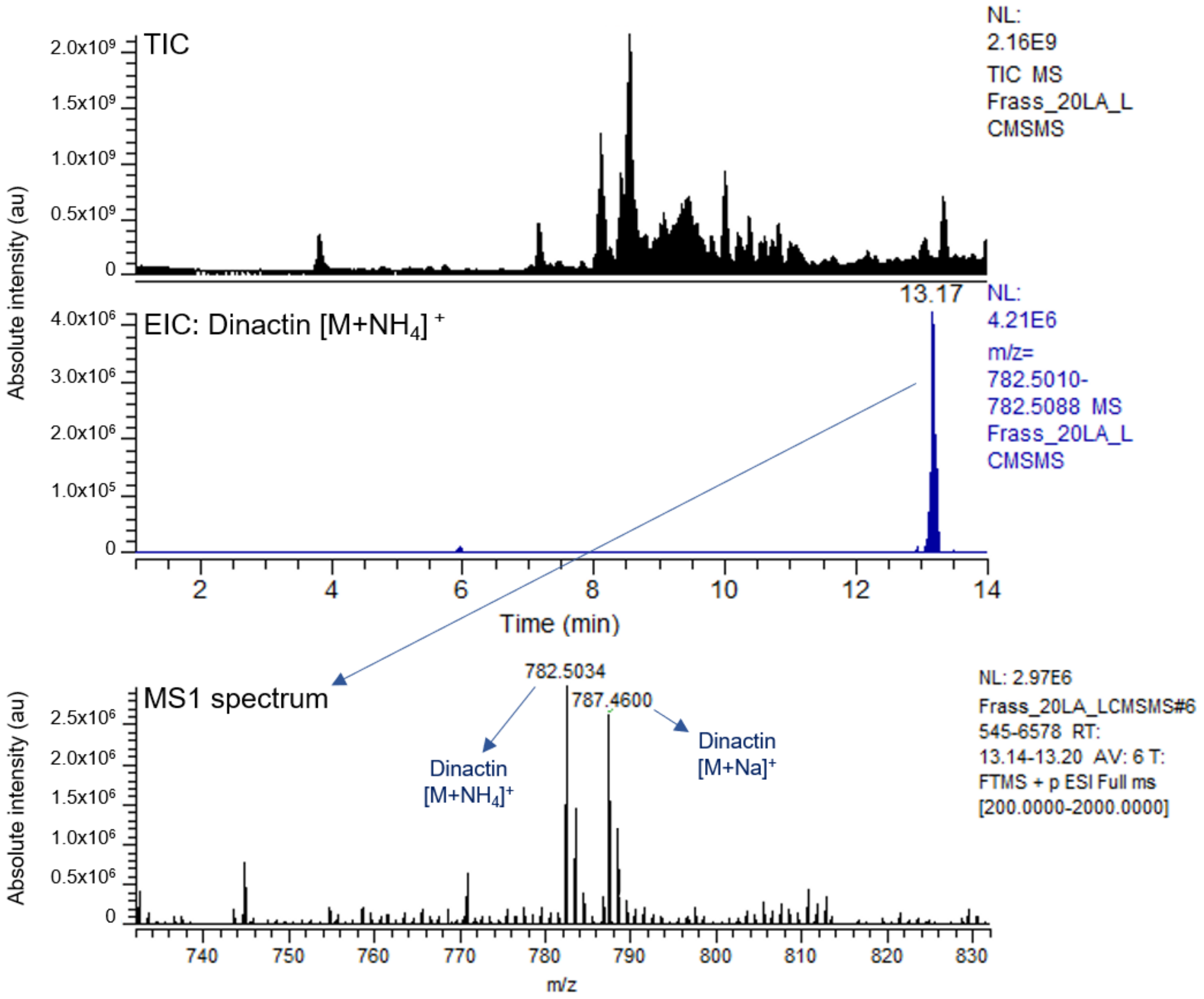


**Appendix 1 - Figure 4:** Three exemplary total ion chromatograms (TIC) of the LC-MS/MS analysis performed on environmental samples (**A**: pupal chamber material; **B, C:** old frass), plus the extracted ion chromatogram (EIC) of an exemplary compound detected in each sample. The MS1 spectrum refers to the main peak detected on each EIC, highlighting two adducts of each compound.

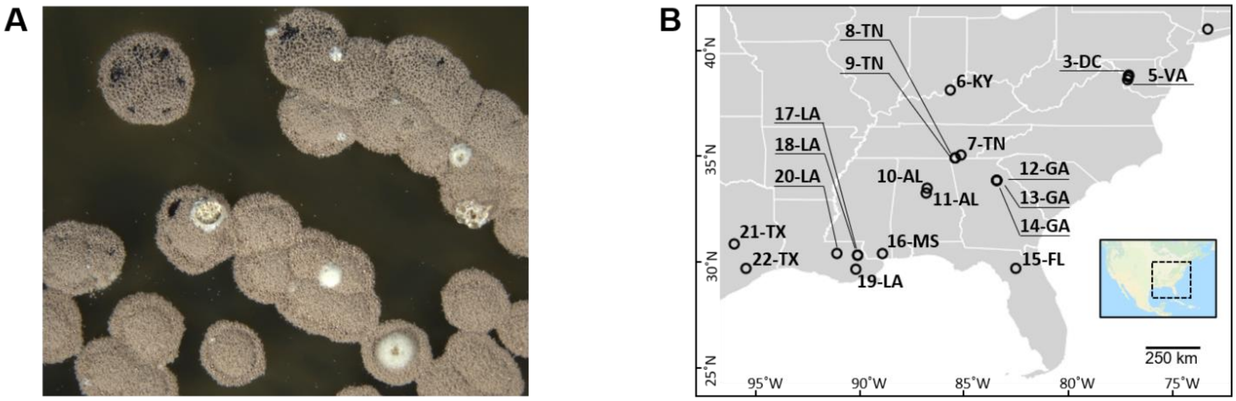


**Appendix 1 - Figure 5:** Strains of *Streptomyces padanus* were isolated from 19/22 of the sampled *O. disjunctus* galleries**. A)** *S. padanus* P333 growing on an ISP2-agar plate after seven days of incubation at 30°C. **B)** Galleries that *S. padanus* was isolated from.

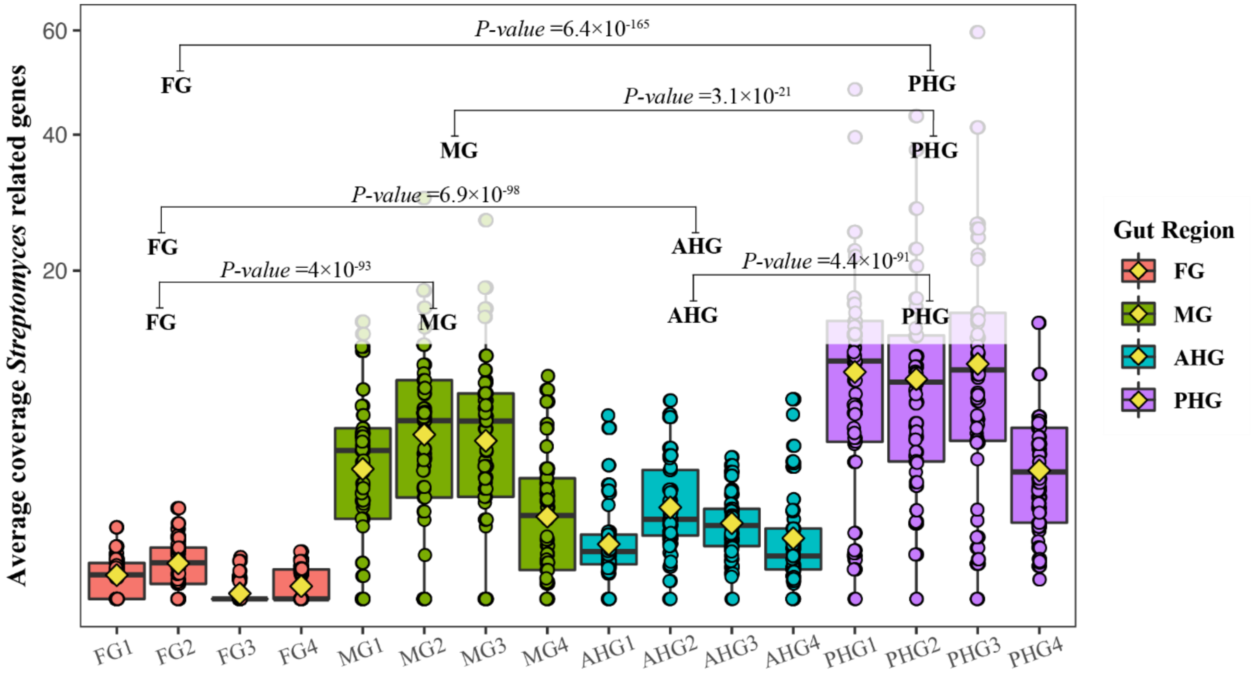


**Appendix 1 - Figure 6:** Average coverage distribution of *Streptomyces*-derived genes identified in the metagenome of the different gut compartments of *O. disjunctus*. The figure shows the prevalence of the *Streptomyces*-derived genes in all four gut compartments with significantly higher normalized-coverage in the posterior hindgut (PHG), the region where frass is compacted prior to its excretion. In each boxplot, a point represents a single gene per category and its detected coverage, and the diamond symbols represent the mean. The box boundaries represent the first and third quartiles of the distribution and the median is represented as the horizontal line inside each box. Boxplots whiskers span 1.5 times the interquartile range of the distribution. FG = Foregut, MG = Midgut, AHG = Anterior Hindgut. Statistical differences were evaluated with Kruskal-Wallis test and pairwise comparisons were done using a two-sided Wilcox test with P-values adjusted using the Benjamini-Hochberg method. Contigs and RPKM-normalized coverage data reported in Ceja-Navarro et al. (2019) (Ceja-Navarro et al., 2019) were used to generate this figure. Contigs were aligned against the NCBI non-redundant database using the DIAMOND software (Buchfink et al., 2014) and the “long reads” option. The obtained alignment was imported into MEGAN (Huson et al., 2018) and the taxonomy assigned using MEGAN’s LCA algorithm for long reads. Coverage data across the four regions of *O. disjunctus*’ gut was retrieved for contigs identified as taxonomically-derived from *Streptomyces* sp.

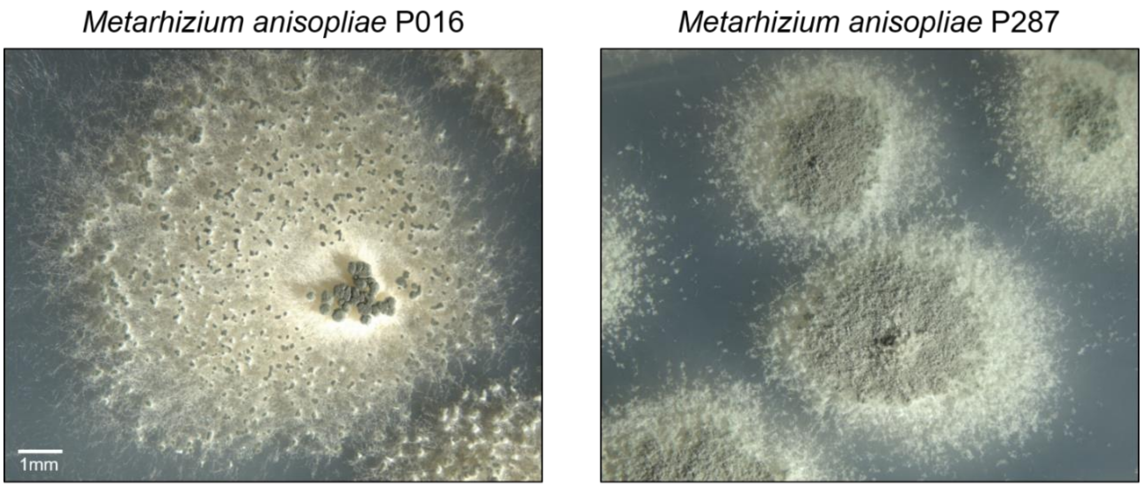


**Appendix 1 - Figure 7:** *Metarhizium anisopliae* strains P016 and P287 phenotypes after 10 days growing on PDA plates incubated at 25°C under constant light. Magnification: 7x.
